# A novel biosensor for ferrous iron developed via CoBiSe: A computational method for rapid biosensor design

**DOI:** 10.1101/2025.07.16.665043

**Authors:** Athanasios Papadopoulos, Manuel T. Anlauf, Jens Reiners, Sueng-Hyun Paik, Aileen Krüger, Benita Lückel, Michael Bott, Thomas Drepper, Julia Frunzke, Holger Gohlke, Stefanie Weidtkamp-Peters, Sander H.J. Smits, Christoph G.W. Gertzen

**Author notes:** Corresponding author Corresponding Author **Sander H.J. Smits** - Center for Structural Studies, Faculty of Mathematics and Natural Sciences, Heinrich Heine University Düsseldorf, Düsseldorf, Germany and Institute for Biochemistry, Faculty of Mathematics and Natural Sciences, Heinrich Heine University Düsseldorf, Düsseldorf, Germany.

## Abstract

Genetically encoded biosensors enable monitoring of metabolite dynamics in living organisms. We present CoBiSe, a computational approach using Constraint Network Analysis to identify optimal insertion sites for reporter modules in molecular recognition elements (MREs). Applied to the iron-binding protein DtxR from *Corynebacterium glutamicum*, CoBiSe identified a flexible connective loop (residues 138-150) for inserting the reporter module, resulting in IronSenseR, a novel ratiometric biosensor for ferrous iron (Fe²⁺). IronSenseR demonstrates high specificity for Fe²⁺ with dissociation constants of 1.55 ± 0.08 µM (FeSO_4_) and 2.44 ± 0.28 µM (FeCl_2_), while showing no binding to Fe³⁺ and other divalent cations. *In vivo* assessment in *Escherichia coli*, *Pseudomonas putida* and *Corynebacterium glutamicum* confirmed IronSenseR’s capability to detect changes in the intracellular iron pool. The creation of IronSenseR underlines that, by reducing search space and eliminating labor-intensive screening, CoBiSe streamlines biosensor development and enables precise creation of next-generation biosensors for diverse metabolites.

## Introduction

Genetically encoded biosensors are minimally invasive tools that allow to monitor changes in various metabolite concentrations in living systems. Functional biosensors combine a molecular recognition element (MRE), which selectively binds a target metabolite, with a reporter module, composed of one or more fluorescent proteins (FP). Fluctuations in metabolite concentration alter the binding to the MRE, which subsequently results in a modulation of the fluorescence signal of the reporter module that can then be detected^1–3^. This powerful feature enables monitoring of cellular dynamics, and provides insights into complex metabolic processes with high spatial and temporal resolution^4^. The design of genetically encoded biosensors has traditionally relied on empirical approaches, where biosensors were created through trial-and-error processes^1, 4, 5^.

Typically, two biosensor designs are favored: The FRET-based biosensors rely on changes in FRET (Förster resonance energy transfer) between two FP domains, whereas the single-FP biosensors are based on environmental modulation of the fluorescence properties of the reporter FP^1, 6^. In both biosensor designs, conformational changes of the MRE upon metabolite binding result in structural rearrangements of the whole biosensor protein and alter the fluorescence properties of the reporter module enabling biosensing readout^1, 4, 6, 7^. While FRET-based biosensors are rather simple to design by sandwiching an MRE in between a suitable donor and acceptor pair of FPs, the creation of single-FP biosensors often requires a more sophisticated split of the MRE to insert the reporter module, which in turn allows to transmit the signal between both biosensor elements^1, 7^. Commonly, single-FP biosensors utilize reporter modules that consist of at least a circular-permutated fluorophore (cpFP), which is fused into the MRE in a peculiar fashion^7^. This fusion of the reporter module to the MRE allows for an environmental-sensitive and conformational-dependent modulation of the fluorescence properties upon metabolite binding to the MRE^7, 8^. Certainly, both popular biosensor designs, FRET-based and single-FP, present advantages and disadvantages: FRET-based biosensors are simple to design and can be used in a ratiometric fashion, but possess a comparably low signal-to-noise ratio, dynamic range and sensitivity^4, 6^. Single-FP biosensors overcome these limitations but support solely intensiometric measurement applications and are more challenging to create^6, 7^. To combine the advantages of both biosensor designs, the Matryoshka biosensor designs was recently developed, which enables the creation of ratiometric biosensors that are excitable at a single wavelength and possess high signal-to-noise ratio, dynamic range, and sensitivity^6, 9, 10^.

While the empirical approach has yielded functional FRET-, single-FP and Matryoshka biosensors in the past, it often requires the labor-intensive and time-consuming creation and screening of large libraries with putative biosensor variants^5, 11, 12^. Additionally, empirical biosensor design approaches may result in biosensors with limited specificity, sensitivity, or dynamic range, and such biosensors frequently require extensive downstream optimization to achieve adequate performance in a biological context^5, 12^. In contrast, previous rational biosensor design approaches leverage detailed knowledge of molecular structures and binding mechanisms of the MRE to systematically engineer biosensors with predictable and tunable properties^7^. These processes are initiated by identifying or designing the biosensor’s modular components, such as the individual MREs and FPs and their linking amino acid residues^7^. Computational modeling and structural biology allowed researchers to predict and enhance interactions between the MRE and its target metabolite, providing a more efficient way to achieve high specificity and affinity^1^. These predictions are often refined and validated experimentally by protein bioengineering approaches using site-directed mutagenesis and directed evolution to further optimize biosensor performance^1, 5^. Although, previous rational design approaches accelerated the development process and yielded biosensors that are inherently more robust, precise, and adaptable, the initial fusion of the MRE to the respective reporter module remained widely random^1, 3, 5, 7, 13^. Current rational approaches for biosensor creation, which already include structural analysis of the MRE, are mostly limited to probing loop-regions or surface residues to identify suitable insertion sites^14–18^. This reduces the search space but is oblivious to the mechanical function of the MRE, thus maintaining too many putative insertion sites of the reporter module and still resulting in non-functional biosensor variants or affecting the biosensors’ affinity and/or specificity. Therefore, a holistic structure-based approach to identify suitable insertion sites would be a major boost in biosensor development. Our goal was to develop a novel, quick and robust method that allows the *in silico* prediction of insertion sites for the reporter module within the MRE without a loss of binding properties for the target metabolite. Hence, we utilized the Constraint Network Analysis (CNA) approach, which is based on protein rigidity theory^19^ and divides the MRE into flexible and rigid parts via a constraint dilution simulation representing thermal unfolding^20, 21^. Rigid parts, in this context, means that no internal motions can propagate efficiently through the interconnected network of stabilizing interactions, facilitating the transfer of conformational changes or dynamic perturbations across the structure while maintaining the overall rigidity of the region^22, 23^. To not disturb the function of the MRE our novel approach intends to identify flexible regions as insertion sites close to the metabolite binding site or preferably between domains connected by it. Thus, rigid parts that are vital for the function of the MRE should not be disturbed by the by the insertion of the reporter module. Ideally the motion and/or forces created upon structural rearrangements by metabolite binding to the MRE should be still transferred between MRE and FP via their collective motions to create and/or trigger a signal in form of changes of the fluorescence properties of the reporter module. The peculiar connection of MRE and reporter module allows for a dynamic fluorescent readout that correlates to changes in metabolite levels.

In summary, the utilization of CNA identifies flexible structural elements within the MRE and discriminates those from rigid clusters within the protein, enabling a highly effective prediction of putative insertion sites for the fusion with the reporter module for the first time. This novel workflow was named CoBiSe, computational prediction for rapid biosensor design. The advantage of the CoBiSe approach is that it strongly reduces the large number of empirically identified or experimentally determined insertion sites. Thus, aiming to identify suitable insertion sites and eliminating the pool of random insertions bypasses the previously required labor-intensive approaches by far and significantly reduces the duration of the whole biosensor creation process. Thus, CoBiSe essentially enhances biosensor creation processes in comparison to previously utilized approaches. To verify that CoBiSe is suitable for the prediction of insertion sites, prominent pre-existing biosensors were used for retrospective insertion site analysis, and the results of identified insertion sites are in line with the described sites in literature. Finally, CoBiSe was utilized to successfully design a completely novel ratiometric Matryoshka biosensor that senses ferrous iron (Fe²⁺), a metabolite for which genetically encoded fluorescent-based biosensors barely exist^24, 25^.

## Results and Discussion

### A retrospective analysis reveals the robustness of the rational biosensor design approach by CoBiSe

First, we looked carefully at the insertion sites of well described biosensors single-FP and tried to identify a rationale for the position of the fluorescent reporter module in the molecular recognition element (MRE). Here, the focus was on MRE into which reporter modules were inserted, that belong to the family of single fluorophore (FP) biosensors. The list of analyzed sensors comprises HyPer3^26^, RexYFP^27^, PercevalHR^28^, MalB2^29^, Tre-CO4^29^, and QUE7µ^30^. Depending on the availability of their protein structures, either the experimentally determined structure or an AlphaFold model^31, 32^ of the MRE was used for further analysis. The respective structural information of the MREs were subjected to flexibility analysis by the Constraint Network Analysis (CNA) software using ensembles of network topologies and fuzzy noncovalent constraints (ENT^FNC^), which does not require generating a structural ensemble but uses an ensemble of network topologies^20, 21^. According to the results of the CAN analysis, regions of the MRE were selected as suitable insertion sites if they met three criteria: I. They are categorized as flexible at the start of or early in the thermal unfolding simulation by CNA, II. They are at the surface of the MRE as adding the reporter module in the core would disturb the function greatly, and III. They were close to the metabolite binding site or, preferably, they connect two different domains or secondary structure elements, that comprise the metabolite binding site. The latter was deemed more promising as a larger movement is to be expected upon metabolite binding, which then may be transferred to the reporter module. These three steps comprise the CoBiSe workflow. The closer the flexible sites are to the binding site and the earlier they become flexible during the thermal unfolding simulation, the more likely they were deemed insertion points. For the aforementioned pre-existing biosensors, the CoBiSe approach resulted in predicted insertion sites in the MRE matching the reported insertion sites in the literature (Figure 1 and Figure S1). Details about the predicted insertion sites can be found in the SI. This poses the question of how much work the application of CoBiSe reduces upon biosensor creation process.

**Figure 1:**
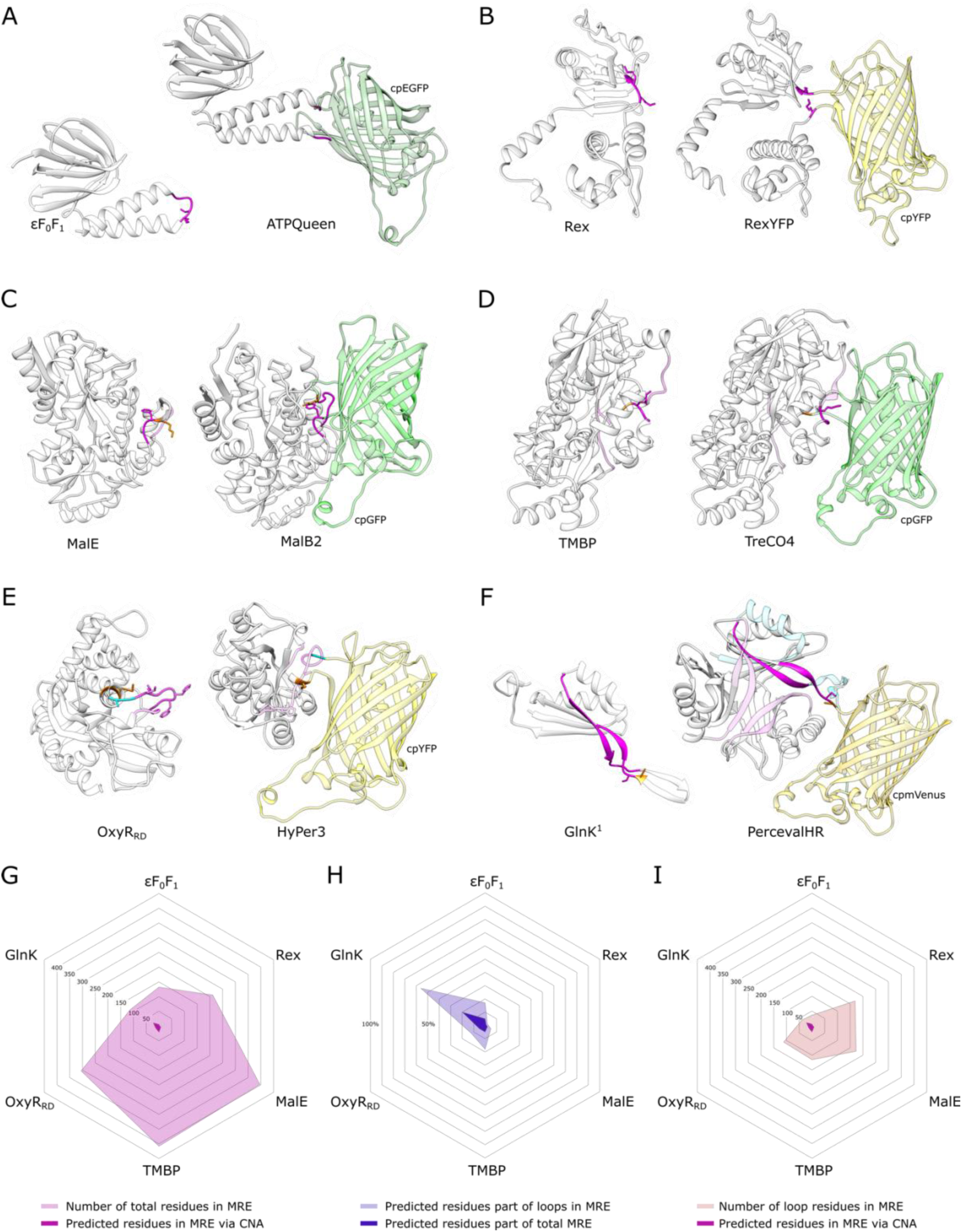
Computational determination of insertion sites for biosensor cassettes. The retrospective computational analysis indicates the utilized insertion sites for the circular-permutated fluorophores in pre-existing biosensors. The AlphaFold3 (AF3) models of the substrate binding protein monomer (left) and the respective biosensor (right) are depicted. The computationally suggested insertion areas (magenta) are highlighted. Flanking residues (orange) and novel insertion sites (cyan) are indicated particularly. Utilized fluorophores are labeled and colored according to their fluorescence emission. **A**: The ε subunit of the bacterial F_0_F_1_ ATP-Synthase was utilized to create ATPQueen biosensor for ATP. **B**: The bacterial transcriptional repressor Rex was utilized to create RexYFP biosensor for NAD^+^/NADH ratio. **C**: The bacterial maltose binding protein MalE was utilized to create MalB2 biosensor for maltose. The utilized insertion site is in line with the predicted area between the indicated flexible loop and the flanking amino acid residue. **D**: The bacterial trehalose-maltose binding protein was utilized and modified to create TreCO4 biosensor for trehalose. The computational analysis determined multiple putative insertion areas (light rose), with the utilized site being most prominent. **E**: The regulatory domain of OxyR H_2_O_2_ binding protein (OxyR_RD_) utilized to create HyPer series of biosensors for H_2_O_2_. The predicted insertion sites are in line with the utilized areas for HyPer biosensor creation. Additional mutational screenings in terms of optimization of fluorescence parameters led to the novel insertion sites (orange and cyan) for HyPer3. **F**: The nucleotide binding protein GlnK was utilized to create Perceval biosensor for ATP/ADP ratio. The protein is active as trimer and only one protomer is used for fluorophore insertion, whereby the determined insertion area is in line with the utilized site. The light cyan region in PercevalHR indicates structural elements of the protomer. **G**: The total number of residues of all tested MREs are indicated (pink) and the predicted residues are highlighted (magenta). **H**: The percentage of predicted residues compared to loop regions of the MREs (light blue) and the percentage of predicted residues compared to the total MREs are indicated (blue) showing the reduced amount of screening for putative insertion sites. **I**: The number of residues in the loop regions of all tested MREs are indicated (rose) and the predicted residues are highlighted (magenta).

Comparing CoBiSe to commonly used random site insertion or whole sequence screening, we achieve a reduction of on average 94 % of the search space, whereas comparing it to a structure-aware approach, which only considers loops regions CoBiSe achieves a reduction of the search space of on average 82 % (Figure 1, G-I). Details for each pre-existing sensor can be found in the SI. In summary, CoBiSe successfully identified insertion sites in six out of six MREs and among those proteins identified seven out of nine insertion sites in a retrospective analysis in published biosensors (Figure 1 and S1). Furthermore, the applied computational analysis of insertion sites reduced the search space tremendously, regardless of whether the entire MRE or only the loop regions were analyzed.

To verify the robustness of CoBiSe in a prospective manner we designed a completely novel ratiometric biosensor for ferrous iron (Fe²⁺).

### CoBiSe analysis revealed insertion sites for a reporter module into iron binding protein

By leveraging structural data in terms to find a suitable iron binding protein for biosensor creation, the global iron regulator (<u>D</u>iphtheria to<u>x</u>in regulator protein, DtxR) from *Corynebacterium glutamicum* was identified. DtxR is a ferrous iron binding protein that can serve as the molecular recognition element (MRE) to be utilized for a putative iron biosensor. By applying the CoBiSe approach for DtxR, it was possible to identify putative insertion sites for the reporter module within the protein (Figure 2, A). A flexible connective loop (residues 139-150) was identified as the optimal insertion site for the Matryoshka biosensor cassette, consisting of superfolder circularly permutated GFP (cpsfGFP) and large stokes shift mApple (LSSmApple), which was successfully integrated into the DtxR protein. Following expression in *Escherichia coli* and protein purification, subsequent functional characterization revealed five insertion variants (I138, D141, D147, S148, and G149) displaying robust positive responses to ferrous iron (0-25 µM) with dynamic ranges exceeding 50% (Figure 2, Figure S2). These variants exhibited significant concentration-dependent increases in cpsfGFP fluorescence upon Fe²⁺ binding, with dynamic ranges between 84.4% and 107%, demonstrating their enhanced sensitivity to DtxR structural changes induced by iron binding. For a detailed description, see SI. Next, to assess the dynamic behavior between the five most responsive variants, the signal to reference correlation of the biosensors was evaluated to determine which variant yields the best signal-to-noise ratio (Figure 2 E). Among the tested biosensor variants, the one created by insertion of the reporter module at position G149 of DtxR demonstrated the highest signal to reference correlation, indicating superior signal-to-noise ratio, sensitivity, and responsiveness to ferrous iron (Figure 2 E). These results highlight that the insertion at position G149 created an effective iron biosensor MDtxR _G149_GA. The superior biosensor variant MDtxR _G149_GA was named IronSenseR and utilized for further applications.

**Figure 2:**
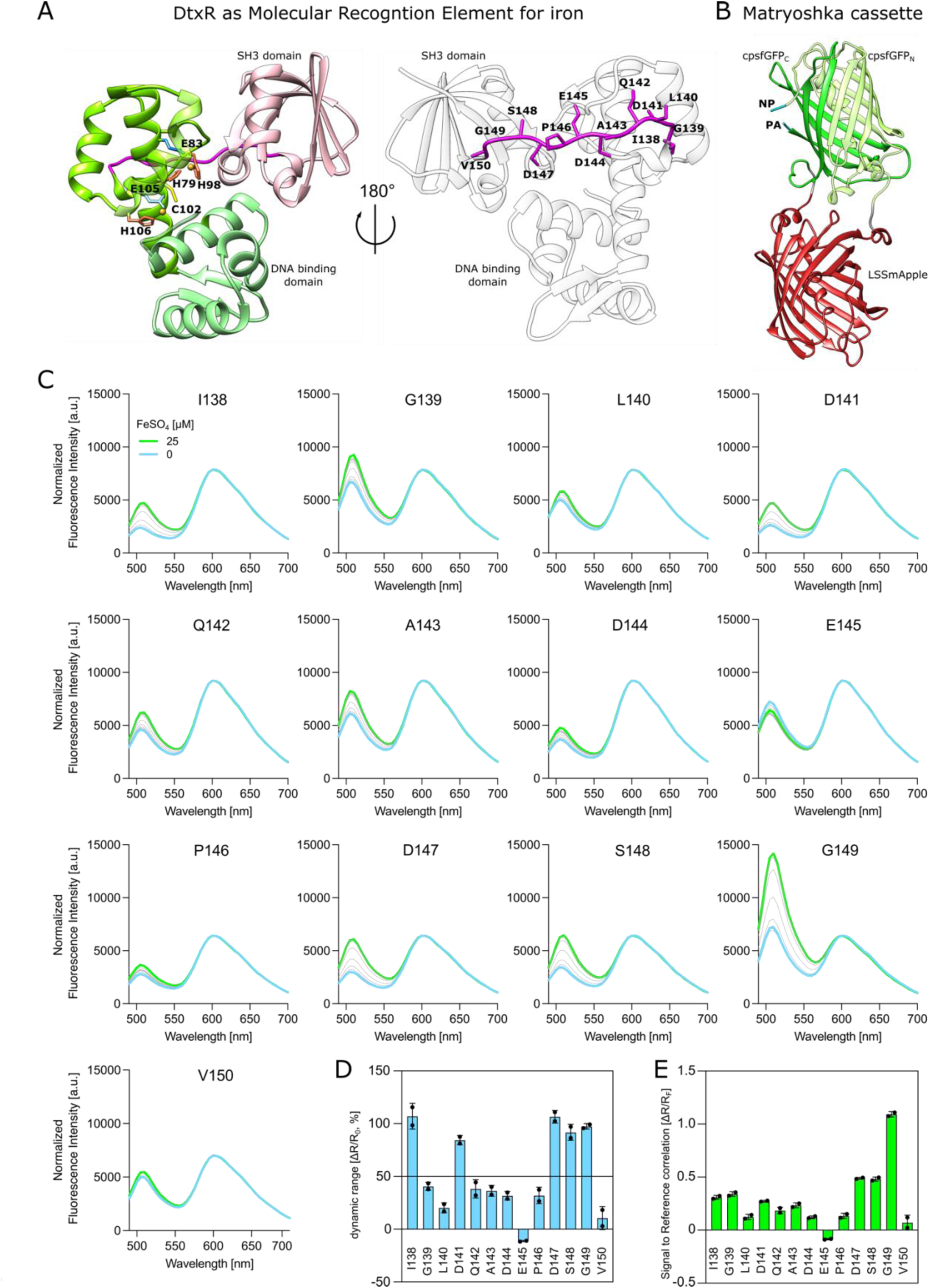
Identification of suitable insertion sites for the creation of responsive biosensors for iron ions. **A**: The AlphaFold3 (AF3) model of DtxR from *C. glutamicum* is depicted. The protein serves as molecular recognition element for iron. The metal binding sites of DtxR (MBS1: H79, E83, H98 and MBS2: E105, C102, H106) and coordinated metal ions are shown. Subdomains (SH3 domain and DNA binding domain) are indicated. The flexible loop (magenta) indicates residues that are used as putative insertion sites for the Matryoshka cassette. **B**: The AlphaFold3 model of the Matryoshka cassette is depicted, the cassette consists of a nested LSSmApple (red) nested to a circular-permutated super folder GFP (cpsfGFP, green). Due to the permutation the amino- and carboxy termini of sfGFP sequence are switched, thus the original C-terminal sequence (sfGFPc, green) appears before the original N-terminal sequence (sfGFP_N_, light green). The linker residues (PA and NP) are indicated. **C**: The titration of FeSO_4_ (0-25 µM) used to monitor the sensing action of DtxR-based ratiometric biosensor variants when insertion of the matryoshka cassette conducted at the proposed positions within the identified flexible loop. **D**: The dynamic range of the respective biosensor variants is depicted [ΔR/R_0_, %]. **E**: The signal-to-reference correlation of the cpsfGFP intensities to the intensity of the reference fluorophore is depicted. The response monitored as an increase of cpsfGFP fluorescence upon ion binding. The variant created upon insertion at position G149 indicates enhanced dynamic range and signal-to-noise ratio in comparison to the other biosensor variants.

### Characterization of the genetically encoded biosensor for ferrous iron

Structural characterization of IronSenseR using SAXS confirmed its monomeric state in solution, with the experimental data matching the AF3 predicted model (χ² = 1.08) ( Figure 3 A, Figure S5, and Table S1). Fluorimetric binding analysis revealed high specificity for ferrous iron (Fe²⁺), with dissociation constants of 1.55 ± 0.08 µM for FeSO₄ and 2.44 ± 0.28 µM for FeCl₂, while showing minimal to no binding with Fe³⁺ compounds and other tested metal ions (MgSO₄, NiSO₄), with only slight responses to MnSO₄ and unsaturated binding to CoSO₄ (Figure 3 B and C, Figure S3 A–H). Control experiments with binding-deficient mutants (H79A, H98A, and C102A) confirmed the specificity of the sensor’s interaction with ferrous iron, as these variants showed negligible fluorescence changes upon Fe²⁺ titration (Figure 3 D, Figure S3 I–K). A detailed characterization can be found in the SI. Together, these results demonstrate IronSenseR’s capability to selectively detect ferrous iron over ferric iron and other divalent cations under in vitro conditions.

**Figure 3:**
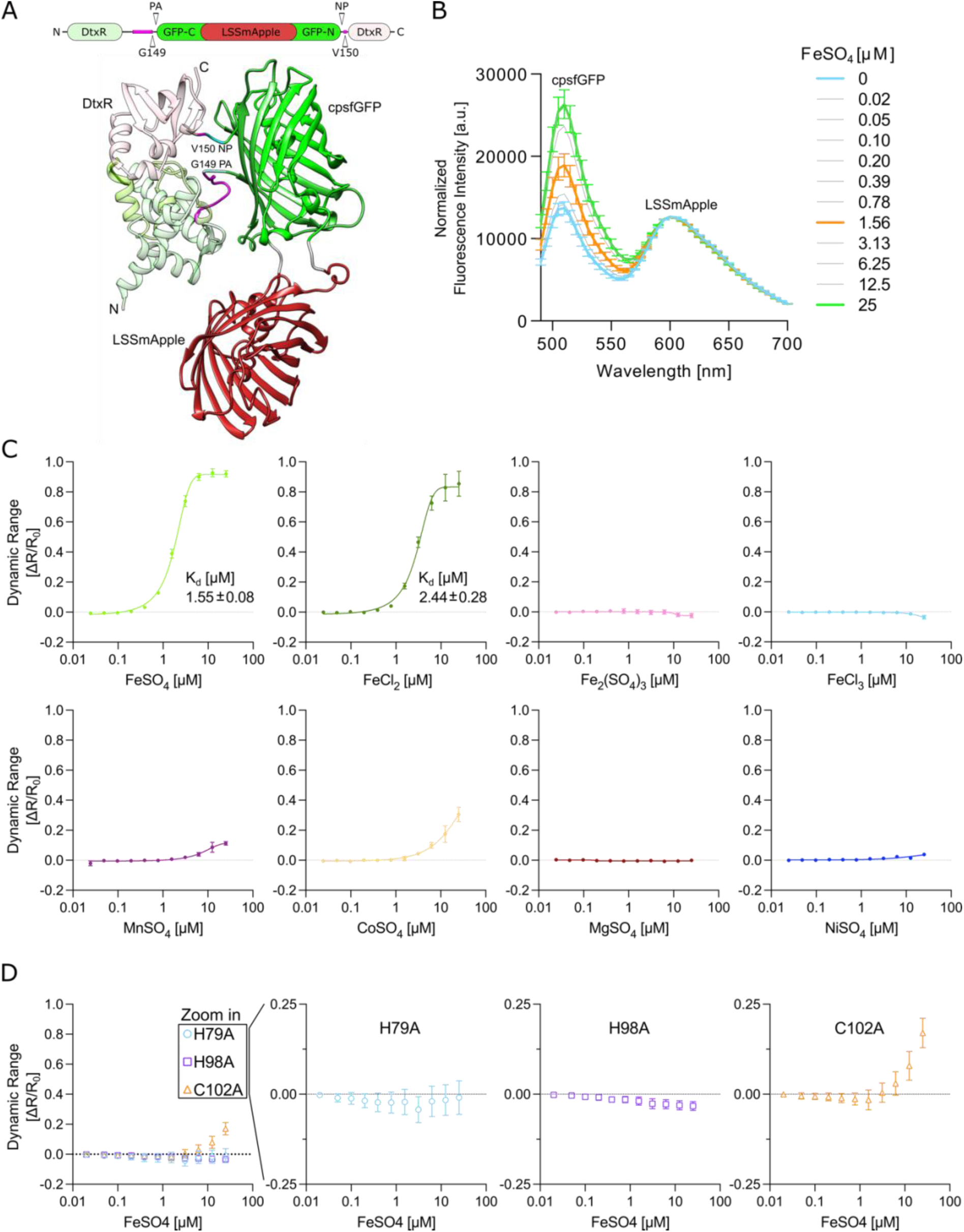
Characterization of IronSenseR: a ratiometric biosensor for ferrous iron. **A**: Molecular architecture and AlphaFold3 (AF3) model of IronSenseR, the Matryoshka biosensor for iron is depicted. The flexible loop (magenta) and the insertion site G149 for the Matryoshka cassette are indicated. **B**: The titration of FeSO_4_ (0-25 µM) used to monitor function of the ratiometric biosensor. **C**: The change of the fluorescence intensities of the reporter FP (cpsfGFP) used to calculate the dynamic range [ΔR/R0] of IronSenseR upon titration of various metal ions at different concentrations. The binding affinities (K_d_’s) are indicated. **D**: The dynamic range for binding-deficient IronSenseR mutants H79A, H98A and C102A are shown. The zoom-in serves for a more detailed view on the data. All the data shown are averages of at least three biological replicates (n=3)

### Assessment of genetically encoded biosensor for ferrous iron *in vivo*

After *in vitro* characterization, IronSenseR was assessed *in vivo* in *Escherichia coli* via fluorescence microscopy. Bacteria were cultivated in nutrient-rich media, and iron availability was subsequently altered after cell growth by adding varying concentrations of the membrane-permeable iron chelator 2,2’-Bipyridine (BPD) ^33, 34^. This was done since iron plays an essential role in the growth and viability of cells, so that alteration of iron homeostasis is challenging and often affects cell growth and protein expression^35^. Microscopy data revealed a reduction in the fluorescence ratio of cpsfGFP/LSSmApple upon increasing BPD concentration (from 1.0 without chelator to 0.84 with 250 µM BPD), with cpsfGFP fluorescence decreasing while LSSmApple fluorescence remained constant (Figure 4 and Figure S4 A-E). Additionally, binding-deficient mutants H79A, H98A and C102A of IronSenseR did not indicate ratiometric changes upon addition of BPD or were greatly diminished in comparison the wild type IronSenseR (Figure 4 and Figure S4 A-E). The ratiometric design allowed analysis of cells with heterogeneous expression levels. While the apparent dynamic range *in vivo* was reduced compared to *in vitro* measurements, likely due to the indirect alteration of intracellular iron and challenges in accurately detecting the change of intracellular iron concentrations, the decreasing cpsfGFP/LSSmApple ratio is in line with the expected sensing ode of IronSenseR. In summary, these results demonstrate that the ratiometric Matryoshka IronSenseR can effectively measure dynamic changes in intracellular iron levels in *E. coli*, though precise correlation between fluorescence ratios and exact or absolute ferrous iron concentrations remains challenging.

**Figure 4:**
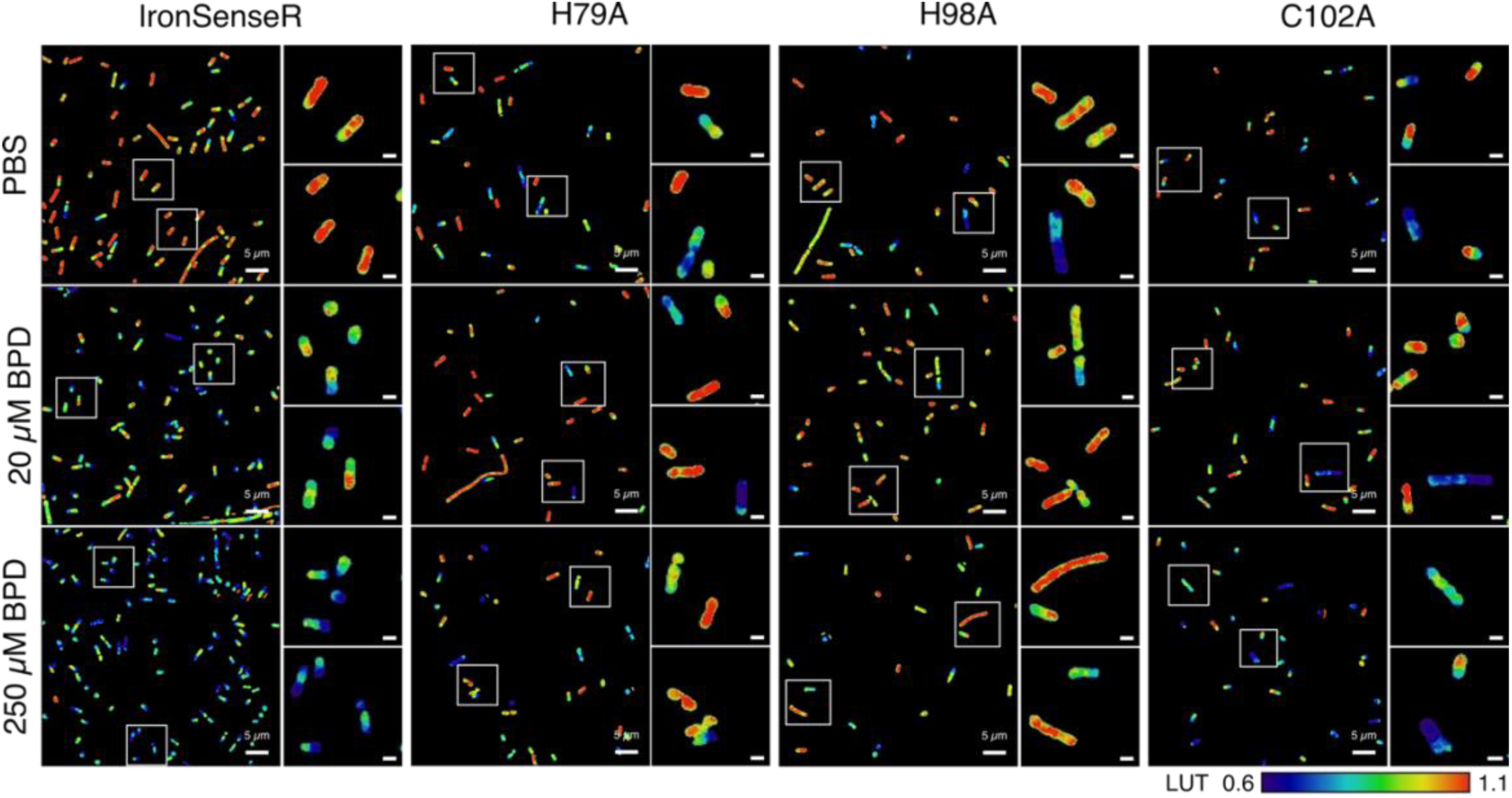
*In vivo* imaging of IronSenseR in *Escherichia coli*. Response of IronSenseR (WT) and binding deficient mutants (H79A, H98A and C102A) to increasing concentrations of the iron chelator 2,2′-Bipyridine (BPD) in bacterial cells. Confocal images of *E. coli* BL21(DE3) expressing the biosensor without addition of BPD (0 µM, PBS top row) and after incubation with either 20 µM BPD (middle row) or 250 µM BPD (bottom row) are depicted. The ratio of green (cpsfGFP) to red fluorescence (LSSmApple) is displayed using a rainbow colored lookup table with two zoomed-in sections depicted in next to the main columns. The scale bar in the zoomed-in images is 1 µm. LUT 0.6-1.1 is depicted by the color bar (down right) indicating the decrease in ratio G/A upon BPD addition. When iron is present, ratio G/A and LUT is high (red) but drops (blue) when iron is chelated by BPD for IronSenseR WT. In case of the binding deficient mutants, this effect is not observable for H79A and H98A or highly diminished as for C102A. Data acquired in biological replicates, n=6 for WT and n=3 for binding deficient mutants.

Next the IronSenseR was utilized in *Pseudomonas putida* and *Corynebacterium glutamicum*, to demonstrate the broad applicability of the biosensor in different bacteria (Figure 5). The corresponding IronSenseR encoding gene-cassette was codon-optimized for its use in *P. putida*, a Gram-negative bacterium relevant to biotechnology and siderophore-based microbial interactions^36, 37^. Under iron depleted conditions, various *Pseudomonas* species produce and secrete the siderophore pyoverdine, which binds environmental ferric iron. The ferri-pyoverdine complex can be specifically taken up by the bacteria, thereby playing an important role in bacterial iron homeostasis^38^. To test if changes in the cytosolic iron pool in *P. putida* can be detected by IronSensR, the biosensor encoding gene-cassette was genomically integrated into the wild-type strain KT2440 (WT) and a pyoverdine-deficient mutant strain (*ΔpvdD*). The recombinant sensor strains were grown under iron-supplemented and iron-depleted conditions. As indicated by the constant fluorescence ratios of the biosensor, increasing the iron concentration in the growth medium did not affect the labile iron pool in the cells of either strain (Figure 5, A–C). However, due to the loss of siderophore-mediated iron acquisition in the *ΔpvdD* strain, increasing concentrations of BPD led to in a detectable decrease in the cytosolic pool of free ferrous iron and a reduction in the fluorescence intensity of the biosensor reporter domain. Conversely, the gradual depletion of iron resulted in the induction of pyoverdine biosynthesis in the WT strain thereby maintaining the intracellular iron pool at a constant level (Figure 5, D–F). These observations confirm the important role of pyoverdine in iron homeostasis and thereby clearly prove the *in vivo* applicability of IronSenseR.

**Figure 5:**
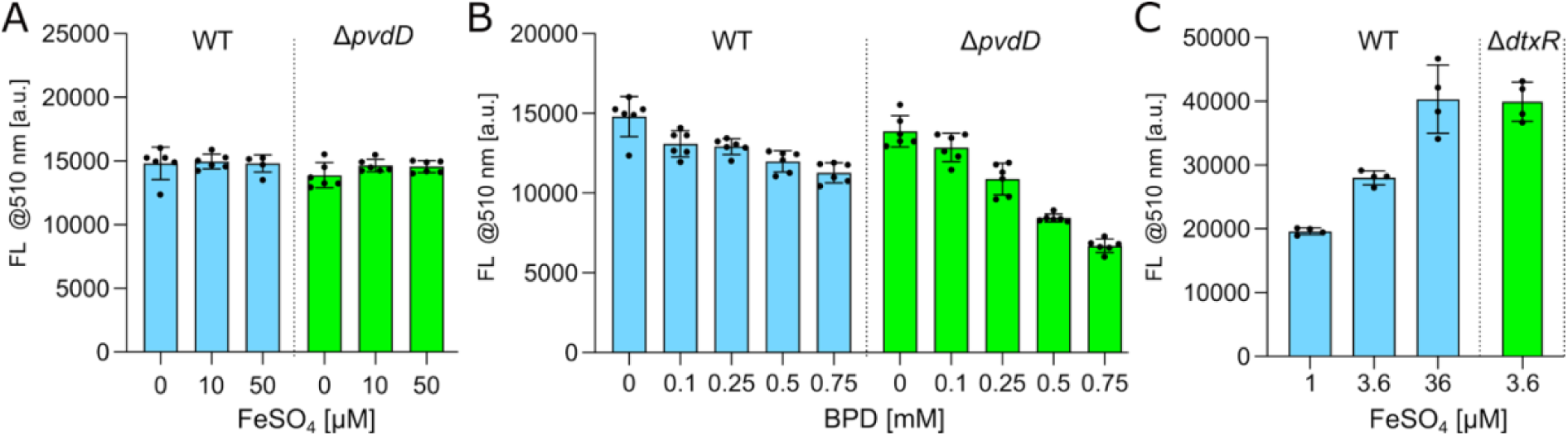
*In vivo* iron sensing in *Pseudomonas putida* and *Corynebacterium glutamicum*. Response of IronSenseR to increasing concentrations of the iron chelator 2,2′-Bipyridine (BPD) and ferrous iron (FeSO_4_) was determined via fluorescence analysis in the *P. putida* wild type (WT) and pyoverdine-lacking mutant (Δ*pvdD*) strain. **A**: Comparison of reporter fluorescence (FL) at 510 nm between *P. putida* WT and Δ*pvdD* upon addition of ferrous iron (FeSO_4_). **B**: Comparison of reporter fluorescence (FL) at 510 nm between *P. putida* WT and Δ*pvdD* upon addition of BPD. **C**: IronSenseR utilized to sense varying iron content upon cultivation of *C. glutamicum*. Increasing iron uptake and/or availability lead to increasing reporter fluorescence of IronSenseR. Comparison of *C. glutamicum* WT and Δ*dtxR* strains upon cultivation in 3.6 µM iron and expression of IronSenseR. The mutant strain Δ*dtxR* lacks the regulation of iron acquisition and is not limited in comparison to the WT, therefore IronSenseR signal is increased indicating a large pool of intracellular iron in comparison to the WT. The Experiments conduct in n=6 biological replicates for *P. putida* and n=4 biological replicates for *C. glutamicum*.

*C. glutamicum* expressing IronSenseR was cultivated with varying iron concentrations from a standard high amount of 36 µM, over a sufficient one of 3.6 µM to a limitation of 1 µM (Figure 5 G). A clear reduction in reporter (cpsfGFP) fluorescence emission signal can be observed upon decreasing iron concentrations. Moreover, a deletion mutant strain (*C. glutamicum*, Δ*dtxR*) was transformed with the IronSenseR encoding plasmid and iron-dependent reporter signals compared to the wild type. DtxR is the master regulator of iron homeostasis in *C. glutamicum*, acting as an iron-activated repressor of iron starvation response^39^. Therefore, it was expected that the mutant accumulates and maintains relatively more intracellular iron than the WT, due to further iron uptake despite iron sufficiency. Using IronSenseR, the expected effect could be observed at the 3.6 µM iron condition, indicated by an increased reporter emission signal in comparison to the wild type (Figure 5 H). Additionally, it is crucial to notify that the Δ*dtxR* mutant possess a growth defect due to iron overload ^40^. This growth defect was indeed observed upon monitoring the optical density (OD600) upon cultivation, indicating that the DtxR-based IronSenseR does not regulatory interfere in *C. glutamicum* with the global regulator pathway.

## Conclusions

The development of biosensors has evolved significantly, transitioning from traditional empirical design methods to rational, computationally informed strategies^1^. Empirical design, while foundational, relies heavily on trial-and-error approaches that are time-consuming, inefficient, and often yield suboptimal biosensors requiring extensive refinement. In contrast, rational design leverages molecular knowledge and computational tools to systematically engineer biosensors with predictable and tunable properties^1, 7^. Constraint Network Analysis (CNA) facilitates the identification of flexible structural elements^20, 21^ within molecular recognition elements (MREs), predicting optimal insertion sites for reporter modules. We demonstrated that our computational approach, CoBiSe, retrospectively predicted insertion sites over a diverse set of MREs (Figure 1) and prospectively led to the successful design of a novel ratiometric Matryoshka biosensor for ferrous iron named IronSenseR.

For the retrospective computational analysis of insertion sites in pre-existing biosensors, we used either high-quality X-ray crystal structures or AlphaFold models^31, 32^ of the MREs. For multimeric MREs, both the multimeric structure and single protomers were analyzed, with both approaches correctly predicting the insertion sites (Figure 1, A–F). This demonstrates the robustness of our approach, showing that no specific structural state or method of structure determination is required. Knowledge of active and inactive protein states is also not a prerequisite, although we are investigating methods to exploit such information to further reduce the search space. Additionally, the ENT^FNC^ approach^21^ implemented in CNA enables this method even when resources for computationally intensive tasks like molecular dynamics simulations are unavailable. Furthermore, MREs with similar folds shared predicted flexible regions (e.g., both MalE in MalB2 and TMBP in Tre-CO4 exhibit a Venus fly trap mechanism), supporting our fundamental concept of viewing MREs as mechanical structures with rigid and flexible parts that move in specific ways to function. This idea is corroborated by investigations showing strong correlations between protein structure and dynamics^41^. Ideally, function should remain undisturbed, so we aim to insert reporter modules in flexible parts less likely to participate in the MRE’s concerted mechanical movements. Insertion sites are not exclusively found in loop regions (though there is a preference), making the CNA flexibility prediction advantageous. Both predicted and experimentally identified sites predominantly occur at transitions between flexible and rigid regions, supporting our hypothesis that when rigid parts move in concert, maximum force is exerted at these transitions. Thus, our CoBiSe approach eliminates the need for labor-intensive random mutagenesis^1, 5^, accelerating development while ensuring optimal sensor performance.

To validate CoBiSe’s capability to predict suitable insertion sites for de novo biosensor creation, we applied CNA to the DtxR protein from *C. glutamicum* to predict insertion sites for a ferrous iron (Fe²⁺) biosensor (Figure 2, A). Ferrous iron plays central roles in numerous cellular processes, serving as a cofactor in oxygen transport, redox reactions, energy production, and gene regulation for bacteria as well as in general for living cells^35, 42, 43^. Its chemical reactivity promotes the generation of damaging reactive oxygen species (ROS)^44^, necessitating tight cellular regulation to balance essential functions against potential toxicity^44^. Previous attempts to create ferrous iron biosensors resulted in limited or indirect detection methods^24, 25^. Our CNA identified a flexible connective loop between I138 and V150 of DtxR as putative insertion sites for the reporter module (Figure 2, A). Inserting the next-generation Matryoshka cassette^9^ (Figure 2, B) into these positions yielded 13 ratiometric biosensor candidates that were rapidly screened for ferrous iron binding in vitro (Figure 2, C). The variant with insertion at position G149 demonstrated superior sensing behavior and high specificity for Fe²⁺ in the physiological micromolar range, without binding to other relevant cations such as Fe³⁺ (Figure 2, C and Figure 3). The next-generation Matryoshka biosensor module provided enhanced signal-to-noise ratio, allowing finely tuned responses upon iron binding. This novel ratiometric biosensor, IronSenseR, was successfully used for *in vivo* analysis of the intracellular iron pool in different bacterial cells (Figure 4 and 5), addressing a critical need for precise and selective detection of ferrous iron in biological contexts^24, 25^. IronSenseR development validates our computational approach, which enabled rapid biosensor creation with optimized dynamic range, specificity, and sensitivity in a single step, demonstrating the adaptability of our design framework.

IronSenseR represents a significant advancement with potential applications in studying iron homeostasis and its dysregulation in cells. Beyond basic research, such tools can support diagnostic screenings of agents targeting iron-related functions in bacterial pathogens^43^. By providing precise, reliable methods to explore biochemical pathways and cellular processes, this technology supports advancements across various research areas.

CoBiSe transforms rational biosensor design by strategically combining structural and computational information, making it adaptable to virtually any protein of interest. This approach enables rapid biosensor creation regardless of target metabolite, as demonstrated by IronSenseR. By significantly narrowing the search space for biosensor cassette insertion sites, CoBiSe offers an efficient, rational, and more economically and environmentally sustainable design process. Its broadly compatible computational requirements ensure accessibility across the scientific community.

## Experimental Section

### Computational insertion site identification

For the application of the Constraint Network Analysis (CNA)^20, 21, 45^ approach in the retrospective analysis on the molecular recognition elements (MREs) of the single fluorophore (FP) biosensors, X-ray crystal structures, AlphaFold models^31, 32^ or available models from the UniProtKB^46^ were used. Here, a deliberate mix of experimentally determined crystal structures and AlphaFold models was used, although an emphasis was put on crystal structures with a high sequence coverage and low resolution. For HyPer3, PercevalHR, and MalB2 AlphaFold models were used and for 30Rex, QUE7µ, and Tre-CO4 the PDBs entry 2DT5^47^, 2E5Y^48^, and 1EU8^49^ were used, respectively. The structures were prepared using the protein preparation wizard in Maestro and protonated to a pH of 7.4 using PROPKA^50^. Subsequently, the structures were analyzed using the CAN approach^45, 51^. For analyzing the rigid cluster decomposition of all MREs, a constraint dilution simulation was performed using CNA on an ensemble of network topologies generated via fuzzy noncovalent constraints (ENT^FNC^)^20, 21^. Subsequently, the unfolding trajectory was visually inspected using VisualCNA^45^ for regions that were determined to be flexible, preferably at the beginning of the thermal unfolding simulation, and that are either near to the metabolite binding site or, preferably, that connect rigid domains that are addressed by the metabolite. VisualCNA is an easy-to-use PyMOL^52^ plugin that allows setting up CNA runs and analyzing CNA results, linking data plots with molecular graphic representations^45^. For the generation of the novel IronSenseR, an AlphaFold model of the DtxR iron-binding protein originating from *Corynebacterium glutamicum* strain (Uniprot ID: Q8NP95) was treated as described above and utilized as MRE for subsequent investigations.

### Molecular cloning

The *dtxR* gene was amplified by PCR from gDNA of *Corynebacterium glutamicum* strain ATCC13032 and sub-cloned into pRSET_B_ based vector and subsequently the correctness was verified by sequencing (Microsynth Seqlab). This vector was linearized by PCR to be used for Gibson assembly. The Matryoshka cassette^9^, encoding for the circular permutated superfolder GFP^53^ (cpsfGFP) and nested large stokes shift mApple (LSSmApple)^9^, was amplified by PCR with corresponding primers possessing suitable ends for Gibson assembly and inserted into the respective insertion sites of DtxR (I138-V150). After PCR, samples were treated with DpnI (NEB) and analyzed by agarose gel electrophoresis. The corresponding amplicons were isolated by gel extraction (MN, gel clean up kit) and utilized for Gibson assembly (NEB). Binding deficient mutants of the biosensor were created by site-directed mutagenesis using KLD mix (NEB). The cloning constructs were used to transform chemically competent *Escherichia coli* DH5α (NEB) by heat shock method. Positive clones were obtained upon cultivation of LB (lysogeny broth, Luria/Miller, Roth) agar plates containing 100 µg/ml ampicillin. Single colonies were used for cultivation in LB supplied with the same antibiotics and were utilized for plasmid isolation (MN Plasmid isolation kit). The plasmids were subjected to DNA sequencing (Microsynth) to confirm successful cloning. Primers for DNA amplification were conducted with corresponding primers listed in supplementary information (Table S2). All kits and protocols followed the manufacturers guidelines.

To apply IronSensR *in vivo* in *Pseudomonas putida*, the corresponding encoding gene-cassette of IronSenseR was optimized using the galaxy codon harmonizer^54^. The resulting biosensor gene was cloned via Gibson assembly^55^ into a miniTn7 vector under the control of a P**_tac_** promoter regulated by the *lac* repressor LacI^56^. To analyze if the loss of siderophore production results in a detectable change of the intracellular pool of free iron ions, the pyoverdine biosynthesis gene *pvdD* was deleted using the pQure system^57^. Using tri-parental conjugation the biosensor expression cassette located on plasmid (pAZ191_GA) was transferred to *P. putida* and subsequently integrated into the genome (Tn7 insertion site) of both the *P. putida* WT strain KT2440 and the *ΔpvdD,* as described previously^58^.

For the *in vivo* usage of IronSenseR in *Corynebacterium glutamicum*, the plasmid vector was exchanged to the shuttle vector pPREx2^59^ , including an IPTG-inducible *tac*-promoter and a kanamycin resistance cassette, while protein-tags were removed. Molecular methods were performed according to standard protocols ^60^. Plasmids were enzymatically assembled using Gibson Assembly ^55^, resulting in pPREx2-MDtxR_G149_GA amplified and stored in *E. coli* DH5α.

### Expression and purification

Biosensor encoding gene cassettes were expressed in *Escherichia coli* BL21 (DE3) and the corresponding proteins were purified by affinity chromatography. In brief, each biosensor variant was encoded by an pRSET_B_-based expression system containing a directly fused N-terminal deca-histidine tag. The expression was conducted in *E. coli* BL21(DE3) (NEB) upon the utilization of autoinduction media^61^. Therefore, chemically competent bacteria were transformed with the respective plasmids by heat shock method and single colonies plated for selective growth on LB agar (lysogeny broth, Luria/Miller, Roth) containing ampicillin (amp, 100 µg/ml) at 37°C for 17 h. A single colony was utilized for expression and used for the inoculation of a pre-culture of 5 ml LB containing ampicillin (100µg/ml). The pre-cultures were cultivated at 37°C, 220 rpm under darkened conditions and the OD_600_ was monitored. Upon reaching OD_600_ of ∼ 0.6, 1.25 ml of pre-cultures were used for inoculation of 50 ml of LB media supplemented with ampicillin (100µg/ml) and containing 0.05% (w/v) glucose as well as 0.2% (w/v) lactose, and put for cultivation at 21°C, 220 rpm for 48 h. After expression, cells were harvested by centrifugation at 4000 x g, 4°C for 40 min. The supernatant was discarded, and the cell pellets were resuspended with 25 ml ice-cold 20mM MOPS pH 7.0 and stored on ice in darkened conditions. An additional centrifugation step at 4000 x g, 4°C for 20 min allowed to remove excess of the supernatant solution, thereby removing putative remaining media compounds. The pellets were again resuspended in 15 ml of 20 mM MOPS pH 7.0 and flash-frozen in liquid nitrogen prior to storage at -80°C.

For purification of the expressed biosensors, the cells were thawed on ice in darkened conditions and 5 ml of the cell suspension was used for cell lysis. Therefore, the cells were pelleted by centrifugation at 11000 x g, 4°C for 1 min and suspended in 2 ml 20 mM MOPS pH 7.0. The cell lysis was conducted by sonification for 2 rounds (Qsonica sonicators) using an amplitude of 50 and 45 pulse cycles with 3 seconds pulse-ON and 8 seconds pulse-OFF. After sonication, the cell lysates were clarified by centrifugation for 20 min at 20830 x g, 4°C to remove cell debris. The clarified lysate containing the histidine-tagged biosensors was applied to NiNTA-based affinity chromatography (Protino, Macherey-Nagel). Therefore, 750 µl NiNTA beads were applied to a gravity flow column (Poly-Prep, Bio-Rad) and rinsed with a total volume of 30 ml deionized water. Subsequently, the beads were equilibrated with 5 ml of 20 mM MOPS pH 7.0 prior to the application of the clarified lysate for immobilization of the histidine-tagged biosensors by gravity flow. The loaded NiNTA beads were washed with 8 ml of 20 mM MOPS pH 7.0, 500 mM KCl and 20 mM imidazole to remove weakly bound impurities. To elute the immobilized biosensors from the NiNTA beads, 3 ml of 20 mM MOPS pH 7.0, 300 mM imidazole were applied, and elution was collected in two 1.5 ml fractions. The buffer of eluted protein was exchanged upon utilization of desalting columns (Cytiva) or size exclusion chromatography via gel filtration on Superdex 200 Increase 10/300 GL column (Cytiva), suiting further structural and biochemical characterization of the biosensor. The protein concentration was determined by UV/Vis spectrometry (NanoDrop, Thermo Fisher Scientific). The purity of the elution fractions was analyzed by SDS-PAGE by conventional Coomassie staining and prior in-gel fluorescence (λ_ex_ 460 nm and λ_em_ 525 nm, Amersham imageQuant 800, GE/Cytiva). The samples were stored for maturation at 4°C for at least 24 h, prior further usage for biochemical, biophysical or structural analysis.

### Functional characterization

The screenings for optimal biochemical conditions of the Matryoshka biosensors for iron were inspired by previous studies^61^. Here, MOPS-based buffer systems were utilized^61^. Fluorescence emission spectra of the biosensors were analyzed on multimode microplate reader (Infinite M Plex, Tecan) inspired by previous protocols ^9, 10, 61^. Therefore, the purified biosensors were diluted to 0.1-0.2 mg/ml in 20 mM MOPS pH 7.0. To assess the binding response of the biosensors to various divalent cations, titrations were conducted in micro titer plates (96 WP, flat bottom, Greiner). Therefore, 50 mM stock solutions of the corresponding ions were generated in deionized water and utilized for subsequent stepwise dilutions to reach 50 µM stocks in 20 mM MOPS pH 7.0. Therefore, 200 µl of the 50 µM stock solution were added to well of lane 12 of each row (A-H) of the micro titer plate, and in all other wells 100 µl of 20 mM MOPS pH 7.0 were added. A serial dilution was performed reaching from 50 to 0.04 µM (from wells 12 to 2) using a multi-channel pipette. Lan1 1 contained only 100 µl of 20 mM MOPS pH 7.0 and equals the 0 control. Afterwards, 100 µl of the corresponding biosensor solution were added to each row of the 96-WP using a multi-channel pipette. An incubation for 15 min at ambient temperature under darkened conditions ensured binding of the ions to the biosensor. The steady-state fluorescence spectra were recorded at 25°C in top reading mode with a bandwidth of 20 nm, 30 flashes, and manual gain of 100 for both, excitation and emission wavelengths. The excitation wavelength (λ_ex_) was 453 nm and the emission spectra were recorded from 490-700 nm in 5 nm steps. The autofluorescence of the buffer was negligibly low. The reporter FP (cpsfGFP) indicated an emission maximum at 505-510 nm and the reference FP at 600 nm. For the data evaluation, the maximal value of the reference FP at 600 nm was used for normalization of each measured data point within the recorded spectra. Furthermore, values of emission maxima were used for calculated relative dynamic range changes (ΔR/R_0_) in response to analyte binding by the respective biosensor as suggested previously ^9, 10^.

To analyze *in vivo* if in *Pseudomonas putida* the loss of siderophore production results in a detectable change of the intracellular pool of free iron ions in pyoverdine biosynthesis deletion background IronSenseR was integrated into the genome) of both the *P. putida* WT strain KT2440 and the *ΔpvdD,* as described previously^58^. The resulting strains including the WT and *ΔpvdD* strains without intergration were pre-cultivated in 1 ml LB at 30° C at 1200 rpm in Flowerplates for 24 hours. For biosensor-based analysis of intracellular iron-levels, the cells were subsequently inoculated in 1 ml LB with an OD_600_ of 0.05. To decrease or increase the iron availability, DIP (0.5 and 7.5 mM) and FeSO_4_ (3.6, 10 and 50 µM) were added to the LB medium, respectively. Cells were cultivated for 4 hours at 30° C (1200 rpm). To induce the expression of the IronSensR gene, 1mM IPTG was added and cell were further incubated for 48 hours at 20° C (1200 rpm).

Electrocompetent *Corynebacterium glutamicum* ATCC13032 WT or Δ*dtxR* cells were transformed with isolated pPREx2-MDtxR_G149_GA plasmid via electroporation^62^. Single colonies (n=4) were cultivated in 5 ml BHI in reaction tubes at 30°C for 5 h. From this pre-culture, a main culture with a starting OD_600_ of 0.01 was inoculated in 15 ml CGXII media containing 2% (w/v) glucose and respective amount of FeSO_4_ as iron source (1 g L^−1^ K_2_HPO_4_, 1 g L^−1^ KH_2_PO_4_, 5 g L^−1^ urea, 42 g L^−1^ MOPS, 13.25 mg L^−1^ CaCl_2_ · 2 H_2_O, 0.25 g L^−1^ MgSO_4_ ·7 H_2_O, 0.27/1/10 mg L^−1^ FeSO_4_ · 7 H_2_O, 10 mg L^−1^ MnSO_4_ · H_2_O, 0.02 mg L^−1^ NiCl_2_ · 6 H_2_O, 0.313 mg L^−1^ CuSO_4_ · 5 H_2_O, 1 mg L^−1^ ZnSO_4_ · 7 H_2_O, 0.2 mg L^−1^ biotin, 30 mg L^−1^ 3,4-dihydroxybenzoate (PCA), 20 g L^−1^ D-glucose, pH 7.0) ^63^ supplemented with kanamycin (25 µg mL^-1^) and 15 µM IPTG in 100 ml shaking flasks on a rotary shaker at 21°C for 48 h. Consequently, cells according to an OD_600_ of 0.5 in 1 ml were harvested via centrifugation (5,000 rpm, 4°C, 5 min) and washed twice with PBS (phosphate buffered saline, 137 mM NaCl, 2.7 mM KCl, 10 mM Na_2_HPO_4_, 1.8 mM KH_2_PO_4_). To ensure full maturation of the reference fluorescence protein LSSmApple, samples were stored overnight in the dark at 4 °C as described previsouly^61^. A volume of 100 µl was analyzed using a multimode microplate reader (Infinite M1000Pro, Tecan) by recording emission spectra from 490 nm to 700 nm in 5 nm increments, with single wavelength excitation at 453 nm and using the maximal value of the reference FP at 600 nm for normalization as described above in the methods section.

### Analysis of biosensor kinetics

To determine the sensing mode of the proposed biosensors and to obtain comparable data, the dynamic range [ΔR/R_0_]^9^, dynamic range in percent [ΔR/R_0_, %] and the Signal to Reference correlation [ΔR/F_R_] were calculated by the following equations:

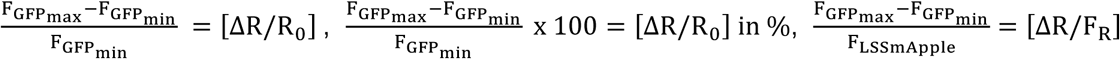

F_GFPmax_: Fluorescence emission of cpsfGFP at 510 nm at highest metabolite concentration

F_GFPmin_: Fluorescence emission of cpsfGFP at 510 nm without metabolite

F_LSSmApple_: Highest emission of the reference FP at 600 nm throughout the assay

The binding affinity was determined by fitting the obtained titration data with the sigmoidal, 4PL, X is log (concentration) equitation of GraphPad Prism 10.4.0 (527) (GraphPad Software, LLC). The representative LogIC50 equals to the Kd values.

### Fluorescence Microscopy

*Escherichia coli* BL21(DE3) were transformed with a plasmid encoding for IronSenseR (MDtxR_G149_GA, WT) or binding deficient mutants of IronSenseR (H79A, H98A and C102A) and cultivated as described above for 24 h in a volume of 20 ml. Prior to imaging, cells were washed with PBS buffer (<u>P</u>hosphate <u>B</u>uffered <u>S</u>aline, 137 mM NaCl, 2.7 mM KCl, 10 mM Na_2_HPO_4_, 1.8 mM KH_2_PO_4_) and incubated for 20 min with PBS buffer containing 0 µM, 20 µM or 250 µM 2,2’-Bipyridine (BPD). Cells were immobilized on Poly-L-Lysine coated 8 Well µ-Slides (ibidi). Imaging was performed using an Olympus Fluoview 3000 confocal laser scanning microscope equipped with a 60x UPLSAPO water objective (NA 1.2). Samples containing IronSenseR were excited with a 488 nm laser (0.5% laser power). Pixel dwell time was set to 2 µs with a line averaging of 2 and a pinhole diameter of 1 AU. Detector range was set to 500-530 nm for cpsfGFP. Fluorescence of LSSmApple was collected at 600-700 nm. Six biological replicates (n=6) for the WT and three biological replicates (n=3) for the binding deficient mutants were analyzed.

### Image processing and image data analysis

For the ratiometric intensity analysis a segmentation was performed using Cellpose 2.0 (Version 2.2)^64^ with the pre-trained “cyto2” model and quantified as well as plotted using a custom python script. In short, the mean fluorescence. Total intensity for the green channel (cpsfGFP, 500-530nm) and red channel (LSSmApple, 600-700 nm) were extracted from the resulting masks and used to calculate the respective ratios. Low and non-expressing cells with an excessively low intensity in either channel were excluded from the analysis using an intensity threshold of 200. This threshold separated best the two peaks (expressing VS non-expressing) of the distribution of individual bacteria mean fluorescence intensities. Fluorescence ratios for all conditions of each biological replicate were normalized to the PBS measurement of the respective replicate. The fluorescence ratios were plotted using a boxplot and a stripplot. For visualization of fluorescence ratios, exemplary images were processed in FIJI^65^. The transmitted light channel was thresholded using either the “Otsu” or “Triangle” method followed by a “Dilate” and an “Open” step. The resulting mask was used for segmentation of fluorescence channels while objects smaller than 0.3 µm² were excluded. A fluorescence ratio channel was created by dividing the pixel intensities of the green channel by the pixel intensities of the red channel. The resulting image was smoothed using a median filter with a radius of 2. Afther that, the normalization factor calculated before was applied to the ratio image. Resulting ratios were displayed using a rainbow-colored lookup table (“physics” in FIJI) ranging from 0.6 to 1.1.

### Structural modelling

The structural models were created upon utilization of AlphaFold 3 prediction algorithm^31, 32^.

### Structural analysis by small angle X-ray scattering

SEC-SAXS data was acquired on beamline BM29 at the ESRF Grenoble^66^. The BM29 beamline was equipped with a PILATUS 2M detector (Dectris) at a fixed distance of 2.827 m. The SEC-SAXS runs were performed at 20°C on a Superdex200 increase 10/300 GL column (100 µl inject, Buffer: 20 mM MOPS pH 7.0, 250 mM KCl) with a flowrate of 0.6 ml/min. We collected 1200 frames with an exposer time of 2 sec/frame. Data were scaled to absolute intensity against water. All used programs for data processing were part of the ATSAS Software package (Version 3.0.5)^67^. Primary data reduction was performed with the programs CHROMIXS ^68^ and PRIMUS ^69^. With the Guinier approximation ^70^, we determine the forward scattering *I(0)* and the radius of gyration (*R_g_*). The program GNOM ^71^ was used to estimate the maximum particle dimension (*D_max_*) with the pair-distribution function *p(r)*. We created a model of MDtxR_G149_GA with AlphaFold3^31, 32^ and compared the theoretical scattering intensity of the resulting models against the experimental data with CRYSOL^72^.

## Associated content

### Supporting Information

The supporting Information is available free of charge at: Link to follow

### Author contributions

AP, CG and SS initiated this study, AP conducted the biosensor design, molecular bioengineering, protein characterization. CG, AP, JR, and HG conducted the computational analysis (CNA and structure prediction) and AP and CG initiated the idea of CoBiSe, MA, TB and SWP conducted the microscopy and imaging evaluation. SP and TD conducted *in vivo* experiments in *P. putida*. AK, BL and JF conducted *in vivo* experiments in *C. glutamicum*. AP, CG and SS supervised the study. AP and SS wrote the initial manuscript. All authors contributed to fruitful discussions and lively development of the manuscript. Correspondence and requests for materials should be addressed to Athanasios Papadopoulos, Sander Smits or Christoph G.W. Gertzen.

### Funding

The research was supported by the German Reseach Foundation (DFG) through the Collaborative Research Center 1535 Microbial Networking (MibiNet, CRC/SFB 1535) Project ID 45809666, (Z01 to SWP and SS, B01 to JF and TD). The Center for Structural Studies (CSS) is funded by DFG, projects 417919780 and INST 208/761-1 FUGG. Center of Advanced imaging is funded by DFG, project 284074525 and project I3D:bio, DFG Grant Number: 462231789.

### Notes

The authors declare no competing financial interest.

## Supporting information

Supplementary infromation

## Acknowledgement

We would like to thank Wolf W. Frommer for inspiring and very valuable discussions and on sharing with us research objectives on Matryoshka biosensor design. Our special thanks to the Institute for Biochemistry for their friendly support in laboratory equipment. Further, we thank the European Synchrotron Radiation Facility for provision of synchrotron radiation facilities, and we would like to thank Dihia Moussaoui for assistance in using beamline BM29. Computational support and infrastructure were provided by the “Zentrum für Informations- und Medientechnologie” (ZIM) at Heinrich Heine University Düsseldorf.

## Abbreviations

MRE, Molecular recognition element

FP, fluorescent protein

FRET, Förster resonance energy transfer

cpFP, circular-permutated fluorophore

sfGFP, super-folder green fluorescent protein

LSS, large stokes shift

Single-FP, single-fluorescent protein or fluorophore

CAN, constraint network analysis

ENT^FNC^, ensembles of network topologies and fuzzy noncovalent constraints

DtxR, Diphtheria toxin regulator protein

BPD, 2,2’-Bipyridine

ROS, reactive oxygen species

## References

1. Frei, M.S., Mehta, S. & Zhang, J. Next-Generation Genetically Encoded Fluorescent Biosensors Illuminate Cell Signaling and Metabolism. Annu Rev Biophys 53, 275–297 (2024).

2. Kim, H., Ju, J., Lee, H.N., Chun, H. & Seong, J. Genetically Encoded Biosensors Based on Fluorescent Proteins. Sensors (Basel) 21 (2021).

3. Wang, M., Da, Y. & Tian, Y. Fluorescent proteins and genetically encoded biosensors. Chem Soc Rev 52, 1189–1214 (2023).

4. Greenwald, E.C., Mehta, S. & Zhang, J. Genetically Encoded Fluorescent Biosensors Illuminate the Spatiotemporal Regulation of Signaling Networks. Chem Rev 118, 11707–11794 (2018).

5. Nadler, D.C., Morgan, S.A., Flamholz, A., Kortright, K.E. & Savage, D.F. Rapid construction of metabolite biosensors using domain-insertion profiling. Nat Commun 7, 12266 (2016).

6. Sadoine, M. et al. Designs, applications, and limitations of genetically encoded fluorescent sensors to explore plant biology. Plant Physiol 187, 485–503 (2021).

7. Nasu, Y., Shen, Y., Kramer, L. & Campbell, R.E. Structure- and mechanism-guided design of single fluorescent protein-based biosensors. Nat Chem Biol 17, 509–518 (2021).

8. Akerboom, J. et al. Crystal structures of the GCaMP calcium sensor reveal the mechanism of fluorescence signal change and aid rational design. J Biol Chem 284, 6455–6464 (2009).

9. Ejike, J.O. et al. A Monochromatically Excitable Green-Red Dual-Fluorophore Fusion Incorporating a New Large Stokes Shift Fluorescent Protein. Biochemistry 63, 171–180 (2024).

10. Ast, C. et al. Ratiometric Matryoshka biosensors from a nested cassette of green- and orange-emitting fluorescent proteins. Nat Commun 8, 431 (2017).

11. Tainaka, K. et al. Design strategies of fluorescent biosensors based on biological macromolecular receptors. Sensors (Basel) 10, 1355–1376 (2010).

12. Koveal, D. et al. A high-throughput multiparameter screen for accelerated development and optimization of soluble genetically encoded fluorescent biosensors. Nat Commun 13, 2919 (2022).

13. Kaczmarski, J.A., Mitchell, J.A., Spence, M.A., Vongsouthi, V. & Jackson, C.J. Structural and evolutionary approaches to the design and optimization of fluorescence-based small molecule biosensors. Curr Opin Struct Biol 57, 31–38 (2019).

14. Ostermeier, M. Engineering allosteric protein switches by domain insertion. Protein Engineering Design and Selection 18, 359–364 (2005).

15. Berrondo, M., Ostermeier, M. & Gray, J.J. Structure prediction of domain insertion proteins from structures of individual domains. Structure 16, 513–527 (2008).

16. Collinet, B. et al. Functionally accepted insertions of proteins within protein domains. J. Biol. Chem. 275, 17428–17433 (2000).

17. Betton, J.-M., Jacob, J.P., Hofnung, M. & Broome-Smith, J.K. Creating a bifunctional protein by insertion of β-lactamase into the maltodextrin-binding protein. Nat. Biotechnol. 15, 1276–1279 (1997).

18. Edwards, W.R., Busse, K., Allemann, R.K. & Jones, D.D. Linking the functions of unrelated proteins using a novel directed evolution domain insertion method. Nucleic Acids Res. 36, e78–e78 (2008).

19. Jacobs, D.J., Rader, A.J., Kuhn, L.A. & Thorpe, M.F. Protein flexibility predictions using graph theory. Proteins 44, 150–165 (2001).

20. Pfleger, C., Rathi, P.C., Klein, D.L., Radestock, S. & Gohlke, H. Constraint Network Analysis (CNA): a Python software package for efficiently linking biomacromolecular structure, flexibility, (thermo-)stability, and function. J Chem Inf Model 53, 1007–1015 (2013).

21. Pfleger, C. & Gohlke, H. Efficient and robust analysis of biomacromolecular flexibility using ensembles of network topologies based on fuzzy noncovalent constraints. Structure 21, 1725–1734 (2013).

22. Thorpe, M.F., Jacobs, D.J., Chubynsky, N. & Rader, A. Generic rigidity of network glasses. Rigidity theory and applications, 239–277 (2002).

23. Gohlke, H. & Thorpe, M.F. A natural coarse graining for simulating large biomolecular motion. Biophys. J. 91, 2115–2120 (2006).

24. Soleja, N. & Mohsin, M. A genetically encoded probe for monitoring and detection of iron in real-time. Sensors & Diagnostics 3, 1714–1723 (2024).

25. Sevimli, G. et al. Probing Subcellular Iron Availability with Genetically Encoded Nitric Oxide Biosensors. Biosensors (Basel) 12 (2022).

26. Bilan, D.S. et al. HyPer-3: a genetically encoded H2O2 probe with improved performance for ratiometric and fluorescence lifetime imaging. ACS Chem. Biol. 8, 535–542 (2013).

27. Bilan, D.S. et al. Genetically encoded fluorescent indicator for imaging NAD+/NADH ratio changes in different cellular compartments. Biochimica et Biophysica Acta (BBA)- General Subjects 1840, 951–957 (2014).

28. Tantama, M., Martínez-François, J.R., Mongeon, R. & Yellen, G. Imaging energy status in live cells with a fluorescent biosensor of the intracellular ATP-to-ADP ratio. Nat. Commun. 4, 2550 (2013).

29. Nadler, D.C., Morgan, S.-A., Flamholz, A., Kortright, K.E. & Savage, D.F. Rapid construction of metabolite biosensors using domain-insertion profiling. Nat. Commun. 7, 12266 (2016).

30. Liang, L. et al. Single-Fluorescence ATP Sensor Based on Fluorescence Resonance Energy Transfer Reveals Role of Antibiotic-Induced ATP Perturbation in Mycobacterial Killing. Msystems 7, e00209–00222 (2022).

31. Jumper, J. et al. Highly accurate protein structure prediction with AlphaFold. Nature 596, 583–589 (2021).

32. Abramson, J. et al. Accurate structure prediction of biomolecular interactions with AlphaFold 3. Nature (2024).

33. Li, S., Crooks, P.A., Wei, X. & de Leon, J. Toxicity of dipyridyl compounds and related compounds. Crit Rev Toxicol 34, 447–460 (2004).

34. Hoyer, J. et al. Proteomic response of Streptococcus pneumoniae to iron limitation. Int J Med Microbiol 308, 713–721 (2018).

35. Andrews, S.C., Robinson, A.K. & Rodriguez-Quinones, F. Bacterial iron homeostasis. FEMS Microbiol Rev 27, 215–237 (2003).

36. Martinez-Garcia, E. & de Lorenzo, V. Pseudomonas putida as a synthetic biology chassis and a metabolic engineering platform. Curr Opin Biotechnol 85, 103025 (2024).

37. Becker, F., Wienand, K., Lechner, M., Frey, E. & Jung, H. Interactions mediated by a public good transiently increase cooperativity in growing Pseudomonas putida metapopulations. Sci Rep 8, 4093 (2018).

38. Ringel, M.T. & Bruser, T. The biosynthesis of pyoverdines. Microb Cell 5, 424–437 (2018).

39. Brune, I. et al. The DtxR protein acting as dual transcriptional regulator directs a global regulatory network involved in iron metabolism of Corynebacterium glutamicum. BMC Genomics 7 (2006).

40. Wennerhold, J. & Bott, M. The DtxR regulon of Corynebacterium glutamicum. J Bacteriol 188, 2907–2918 (2006).

41. Hensen, U. et al. Exploring protein dynamics space: the dynasome as the missing link between protein structure and function. PloS one 7, e33931 (2012).

42. Frawley, E.R. & Fang, F.C. The ins and outs of bacterial iron metabolism. Mol Microbiol 93, 609–616 (2014).

43. Ratledge, C. & Dover, L.G. Iron metabolism in pathogenic bacteria. Annu Rev Microbiol 54, 881–941 (2000).

44. Bradley, J.M. et al. Bacterial iron detoxification at the molecular level. Journal of Biological Chemistry 295, 17602–17623 (2020).

45. Rathi, P.C., Mulnaes, D. & Gohlke, H. VisualCNA: a GUI for interactive constraint network analysis and protein engineering for improving thermostability. Bioinformatics 31, 2394–2396 (2015).

46. UniProt, C. UniProt: a worldwide hub of protein knowledge. Nucleic Acids Res 47, D506–D515 (2019).

47. Nakamura, A., Sosa, A., Komori, H., Kita, A. & Miki, K. Crystal structure of TTHA1657 (AT-rich DNA-binding protein; p25) from Thermus thermophilus HB8 at 2.16 A resolution. Proteins 66, 755–759 (2007).

48. Yagi, H. et al. Structures of the thermophilic F1-ATPase epsilon subunit suggesting ATP-regulated arm motion of its C-terminal domain in F1. Proc Natl Acad Sci U S A 104, 11233–11238 (2007).

49. Diez, J. et al. The crystal structure of a liganded trehalose/maltose-binding protein from the hyperthermophilic Archaeon Thermococcus litoralis at 1.85 A. J Mol Biol 305, 905–915 (2001).

50. Rostkowski, M., Olsson, M.H., Sondergaard, C.R. & Jensen, J.H. Graphical analysis of pH-dependent properties of proteins predicted using PROPKA. BMC Struct Biol 11, 6 (2011).

51. Hensen, U. et al. Exploring protein dynamics space: the dynasome as the missing link between protein structure and function. PLoS One 7, e33931 (2012).

52. Schrödinger, LLC, Edn. Version 2.5 (2022).

53. Pedelacq, J.D., Cabantous, S., Tran, T., Terwilliger, T.C. & Waldo, G.S. Engineering and characterization of a superfolder green fluorescent protein. Nat Biotechnol 24, 79–88 (2006).

54. Claassens, N.J. et al. Improving heterologous membrane protein production in Escherichia coli by combining transcriptional tuning and codon usage algorithms. PLoS One 12, e0184355 (2017).

55. Gibson, D.G. et al. Enzymatic assembly of DNA molecules up to several hundred kilobases. Nat Methods 6, 343–345 (2009).

56. Choi, K.H. & Schweizer, H.P. mini-Tn7 insertion in bacteria with single attTn7 sites: example Pseudomonas aeruginosa. Nat Protoc 1, 153–161 (2006).

57. Volke, D.C., Friis, L., Wirth, N.T., Turlin, J. & Nikel, P.I. Synthetic control of plasmid replication enables target- and self-curing of vectors and expedites genome engineering of Pseudomonas putida. Metab Eng Commun 10, e00126 (2020).

58. Wynands, B. et al. Metabolic engineering of Pseudomonas taiwanensis VLB120 with minimal genomic modifications for high-yield phenol production. Metab Eng 47, 121–133 (2018).

59. Bakkes, P.J. et al. Improved pEKEx2-derived expression vectors for tightly controlled production of recombinant proteins in Corynebacterium glutamicum. Plasmid 112, 102540 (2020).

60 Sambrook, J., Fritsch, E. & Maniatis, T. Molecular cloning: a laboratory manual. Plainview. New York: Cold Spring Harbor Laboratory Press, chapt 16, 66 (1989).

61. Sadoine, M. et al. Affinity Purification of GO-Matryoshka Biosensors from E. coli for Quantitative Ratiometric Fluorescence Analyses. Bio Protoc 10, e3773 (2020).

62. van der Rest, M.E., Lange, C. & Molenaar, D. A heat shock following electroporation induces highly efficient transformation of Corynebacterium glutamicum with xenogeneic plasmid DNA. Appl Microbiol Biotechnol 52, 541–545 (1999).

63. Keilhauer, C., Eggeling, L. & Sahm, H. Isoleucine synthesis in Corynebacterium glutamicum: molecular analysis of the ilvB-ilvN-ilvC operon. J Bacteriol 175, 5595–5603 (1993).

64. Pachitariu, M. & Stringer, C. Cellpose 2.0: how to train your own model. Nat Methods 19, 1634–1641 (2022).

65. Schindelin, J., et al. Fiji: an open-source platform for biological-image analysis. Nat Methods 9, 676–682 (2012).

66. Tully, M.D. et al. BioSAXS at European Synchrotron Radiation Facility - Extremely Brilliant Source: BM29 with an upgraded source, detector, robot, sample environment, data collection and analysis software. Journal of synchrotron radiation 30, 258–266 (2023).

67. Manalastas-Cantos, K. et al. ATSAS 3.0: expanded functionality and new tools for small-angle scattering data analysis. J Appl Crystallogr 54, 343–355 (2021).

68. Panjkovich, A. & Svergun, D.I. CHROMIXS: automatic and interactive analysis of chromatography-coupled small angle X-ray scattering data. Bioinformatics (2017).

69. Konarev, P.V., Volkov, V.V., Sokolova, A.V., Koch, M.H.J. & Svergun, D.I. PRIMUS: a Windows PC-based system for small-angle scattering data analysis. Journal of Applied Crystallography 36, 1277–1282 (2003).

70. Guinier, A. Small-angle X-ray diffraction: application to the study of ultramicroscopic phenomena. Annales de Physique 11, 161–237 (1939).

71. Svergun, D.I. Determination of the Regularization Parameter in Indirect-Transform Methods Using Perceptual Criteria. J Appl Crystallogr 25, 495–503 (1992).

72. Svergun, D., Barberato, C. & Koch, M.H.J. CRYSOL– a Program to Evaluate X-ray Solution Scattering of Biological Macromolecules from Atomic Coordinates. Journal of Applied Crystallography 28, 768–773 (1995).

