## Supplementary infromation for "A novel biosensor for ferrous iron developed via CoBiSe: A computational method for rapid biosensor design"

### corresponding author

#### Suplemental Results

**Predicted insertion sites for retrospective analysis**

For QUE7μ (Figure 1, A), these are residues 106-111 as they are flexible residues that lie between the two helices connected by the binding site, with the reported insertion site being between residue 106-107. For RexYFP (Figure 1, B), these are residues 79-81 as they are flexible residues that lie between two domains close to the binding site, with the reported insertion site between residue 79-80. The MRE of RexYFP can form a homodimer. Applying this method to the homodimer also predicts residues 78-81 as a possible insertion site (Figure S1). For MalB2 (Figure 1, C), these are residues 193-195 and 204-208 as they are flexible residues that lie between the two domains connected via the binding site with the reported insertion site being between residue 195-196. For Tre-CO4 (Figure 1, D), these are residues 95-101, 287-292, 335-337, and 362-366 as they are flexible residues that lie between the two domains connected via the binding site, with the reported insertion site being between residue 334 and 335. Here, due to the higher base rigidity of the MRE, these residues became flexible during step 7 of 136 of the thermal unfolding simulation. For HypPer3 (Figure 1, E) the analysis predicted an insertion site located at positions 187-190 and 213-221 as they are flexible residues that are close to the site where a disulfide bridge is formed in the presence of hydrogen peroxide ($H_2O_2$) with the reported insertion site being at residue 206. After the previous five successes, this prompted us to investigate further into the design of this biosensor. HyPer3 is an improved variant out of the HyPer series for which four possible insertion sites were identified in the MRE of which the one at 206 showed the best ratiometric response, which is why it was chosen for further improvement[1]. The reported insertion sites in the MRE of HyPer and thus, HyPer3 are between 205 and 206, 211 and 212, 214 and 215, and 218 and 219, with the latter two predicted via this approach. Finally, for PercevalHR (Figure 1, F) the residues 26-41 and 54-59 are flexible residues that are close to the binding site, with the reported insertion site being between residue 53-54. Since PercevalHR can appear homotrimeric, the method was also applied to the trimer. Here, the range of residues is even smaller compared to the monomer with residues 37-41 and 54-55 (Figure S1). Thus far, our retrospective insertion site analysis predicted the same insertion sites for the reporter module as they were utilized for the described pre-existing biosensors (Figure 1).

**Reduction in search space compared to common approaches**

Commonly, previous approaches screened for putative insertion sites throughout the whole MRE, thus this is a good starting point to estimate the efficiency of our approach. Considering the whole MREs respectively as possible insertion sites, using this approach means a reduction of the search space of 95% (6 vs. 132) for QUE7μ, 99% (3 vs. 211) for RexYFP, 98% (8 vs. 396 residues) for MalB2, 95% (21 vs. 408) for Tre-CO4, 95% (13 vs. 305) for

HyPer3, and 80% (22 vs. 112) for PercevalHR (Figure 1, G and H). Hence, the application of our rational computer-based approach on average leads to an efficient reduction the search space by 94% when screening for insertion sites within the whole MRE (Figure 1, H). Some biosensor design approaches probe only in the loop regions of the MRE, as possible insertion sites can be identified there more frequently[2]. This reduces the search space so that our approach must also compete against this rationale. If only the loop regions of the MRE are considered as a form of structure-based prediction, the reduction of the search space is 83% (6 vs. 36) for QUE7µ, 98% (3 vs. 169) for RexYFP, 95% (8 vs. 173 residues) for MalB2, 82% (21 vs. 115) for Tre-CO4, 88% (13 vs. 111) for HyPer3, and 44% (22 vs. 39) for PercevalHR (Figure 1, H and I). Of course, this metric is worse than before but here CoBiSe still reduces the search space by 82% on average.

**Design of a biosensor for ferrous iron**

Using CoBiSe, a large connective loop ranging from residues 139 to 150 was identified (Figure 2, A). These residues are potential insertion sites due to their flexibility and position between two domains connected by the proposed loop. This amounts to a reduction in search space by 95% (12 of 230) for the complete MRE and 84% (12 of 74) considering only loop regions. Next, the Matryoshka biosensor cassette[3], consisting of superfolder circularly permutated GFP (cpsfGFP) and large stokes shift mApple (LSSmApple) (Figure 2, B), was successfully integrated in between every amino acid within the identified flexible connective loop of the DtxR protein (Figure 2 A and B). Putative iron biosensors were successfully expressed in bacterial systems, with all designed variants exhibiting detectable expression levels as confirmed by downstream purification procedures (Figure S2). Subsequent purification of these variants was achieved using affinity chromatography, yielding proteins that correspond to the expected molecular weight of approximately 80 kDa, as determined by SDS-PAGE (Figure S2). These results were in line with the expected theoretical MW of 84.19 kDa. The samples appeared pure in the SDS-PAGE (Figure S2, A) and the in-gel fluorescence revealed no other bands suggesting that no major fluorescent degradation products occurred and more importantly no interfering fluorescence signal will arise during further fluorimetric analysis upon titration of metal ions (Figure S2, B). Next, the putative biosensor variants were screened for iron sensing action (Figure 2, C). Functional characterization of the purified biosensors was conducted by fluorimetric analysis screening for their response to iron (II) sulfate ($FeSO_4$) titrations (Figure 2 C). Insertion variants at positions I138, D141, D147, S148, and G149 displayed an obvious ferrous iron ($Fe^{2+}$) concentration dependent increase in cpsfGFP fluorescence compared to their unbound state (Figure 2, C). This indicates a robust and positive responsive configuration for these sites, suggesting enhanced sensitivity to structural changes of DtxR induced by metabolite binding. Conversely, insertion variants at position G139, L140, Q142, A143, D144, P146, and V150 exhibited only moderate changes in fluorescence intensity, suggesting a more subdued responsiveness in these configurations (Figure 2, C). Out of all tested variants, only E145 indicated a negative response correlation of cpsfGFP fluorescence upon ferrous iron binding (Figure 2, C). The dynamic range was employed to facilitate quantitative comparisons among the different biosensor variants by providing a normalized metric (Figure 2, D). Twelve variants indicated an increase in dynamic range, underlining a positive sensing mode of the novel Matryoshka iron biosensors. Here, five variants indicated an enhanced dynamic range above 50%, being I138 with 107%, D141 with 84.4%, D147 with 106.6%, S148 with 91.8%, and G149 with 97.5% (Figure 2, D).

#### Characterization of IronSenseR

The IronSenseR was characterized *in vitro* by biochemical and structural analysis (Figure 3). Structural investigations by small angle X-ray scattering (SAXS) confirmed the monomeric state of the biosensor in solution, furthermore we compared the theoretical scattering intensity of the AF3 predicted models and identified the best-fit model ($\chi2 = 1.08$) with the experimental data (Figure 3, A, Figure S5 and Table S1). IronSenseR indicates an enhanced dynamic range for ferrous iron ($Fe^{2+}$) in form of iron (II) sulfate (Figure 3, B). To assess the biosensors' specificity towards further ions, ferrous iron in form of iron (II) chloride ($FeCl_2$), ferric iron ($Fe^{3+}$) as iron (III) sulfate ($Fe_2(SO_4)_3$) and iron (III) chloride ($FeCl_3$), manganese (II) sulfate ($MnSO_4$), cobalt (II) sulfate ($CoSO_4$), magnesium (II) sulfate ($MgSO_4$), and nickel (II) sulfate ($NiSO_4$) were used for further investigations (Figure 3, C and Figure S3). Fluorimetric binding analysis upon ion titrations revealed the highest affinity for ferrous iron ($Fe^{2+}$) in form of $FeSO_4$, with a dissociation constant ($K_d$) of 1.55 ± 0.08 µM, indicating strong and specific binding (Figure 3, C). Similarly, the sensor showed high affinity for $FeCl_2$, with a $K_d$ of 2.44 ± 0.28 µM, being within a comparable range to $FeSO_4$ (Figure 3 C). In contrast, negligible changes in cpsfGFP fluorescence and dynamic range were observed for $Fe_2(SO_4)_3$, $FeCl_3$, $MgSO_4$, and $NiSO_4$, suggesting minimal to no binding to these compounds (Figure 3 C). A slight response was detected for $MnSO_4$, but it exhibited a significantly reduced dynamic range compared to $FeSO_4$ and $FeCl_2$. Interestingly, $CoSO_4$ displayed titratable binding but did not reach saturation in the applied concentration range, implying lower affinity (Figure 3 C). These findings demonstrate that IronSenseR preferentially binds ferrous iron ($Fe^{2+}$), while showing little to no interaction with ferric iron ($Fe^{3+}$) or other tested metal ions (Figure 3 C). To address whether the obtained data is indeed due to the binding of ferrous iron to the biosensor, binding deficient mutants were created and utilized for fluorimetric analysis upon titration of $FeSO_4$. As described in literature, DtxR contains two metal ion binding sites (MBS 1 and 2)[4]. To create binding deficient biosensors the coordinating residues H79, H98, and C102 were exchanged with alanine [4, 5]. The corresponding mutants of MBS1, H79A and H98A, indicate nearly no binding to ferrous iron (Figure 3, D and Figure S3). Negligible changes in fluorescence were observed for MBS 2 mutant C102A, concluding also no binding to ferrous iron in a relevant concentration range (Figure 3, D). Taken together, the described results demonstrate the capability of IronSenseR to bind selectively ferrous iron. Moreover, the results point to the specificity and the suitability of the biosensor for detecting ferrous iron over ferric iron or other divalent cations under *in vitro* conditions.

### Supplemental Figures

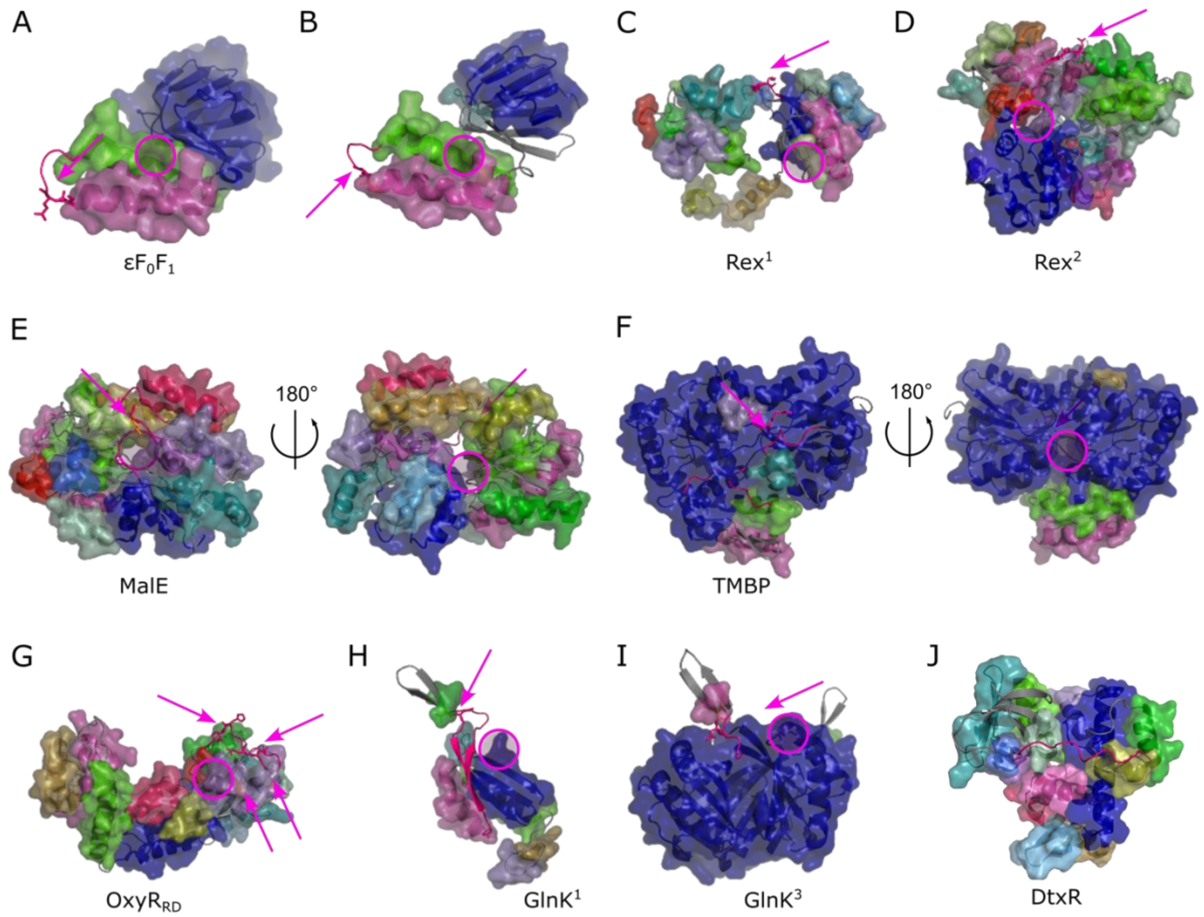


**Figure S1: CNA results with insertion point prediction for the molecular recognition element**

The results of the CNA analysis using $ENT^{FNC}$ for each of the molecular recognition elements of previously published single fluorophore biosensors are shown; for $\varepsilon F_0F_1$ (QUE7µ) (**A**) and an element similar to QUE7µ (**B**) but with a different loop and N-terminal helix sequence and length, for monomeric $Rex^1$ (RexYFP) (**C**) and its homodimeric $Rex^2$ form (**D**), for MalE (MalB2) (**E**) and its 180° rotated view, TMBP (Tre-CO4) (**F**) and its backside view, for $OxyR_{RD}$ (HyPer3) (**G**) for which the sites found in the generation of HyPer1 are shown, for monomeric $GlnK^1$ (PercevalHR) (**H**) and its homotrimeric $GlnK^3$ form (**I**) are shown, (**J**) for DtxR monomer, the metal ion binding protein used for the creaton of IronSenseR is shown. For the predicted insertion site, the cartoon representation is colored magenta with the stick representation indicating the insertion site in the published sensors. If a stick representation is colored orange, it is part of the insertion site, but not predicted. The insertion site is also indicated by a magenta arrow. The (metabolite) binding site in the molecular recognition element is shown as a plum and pink circle.

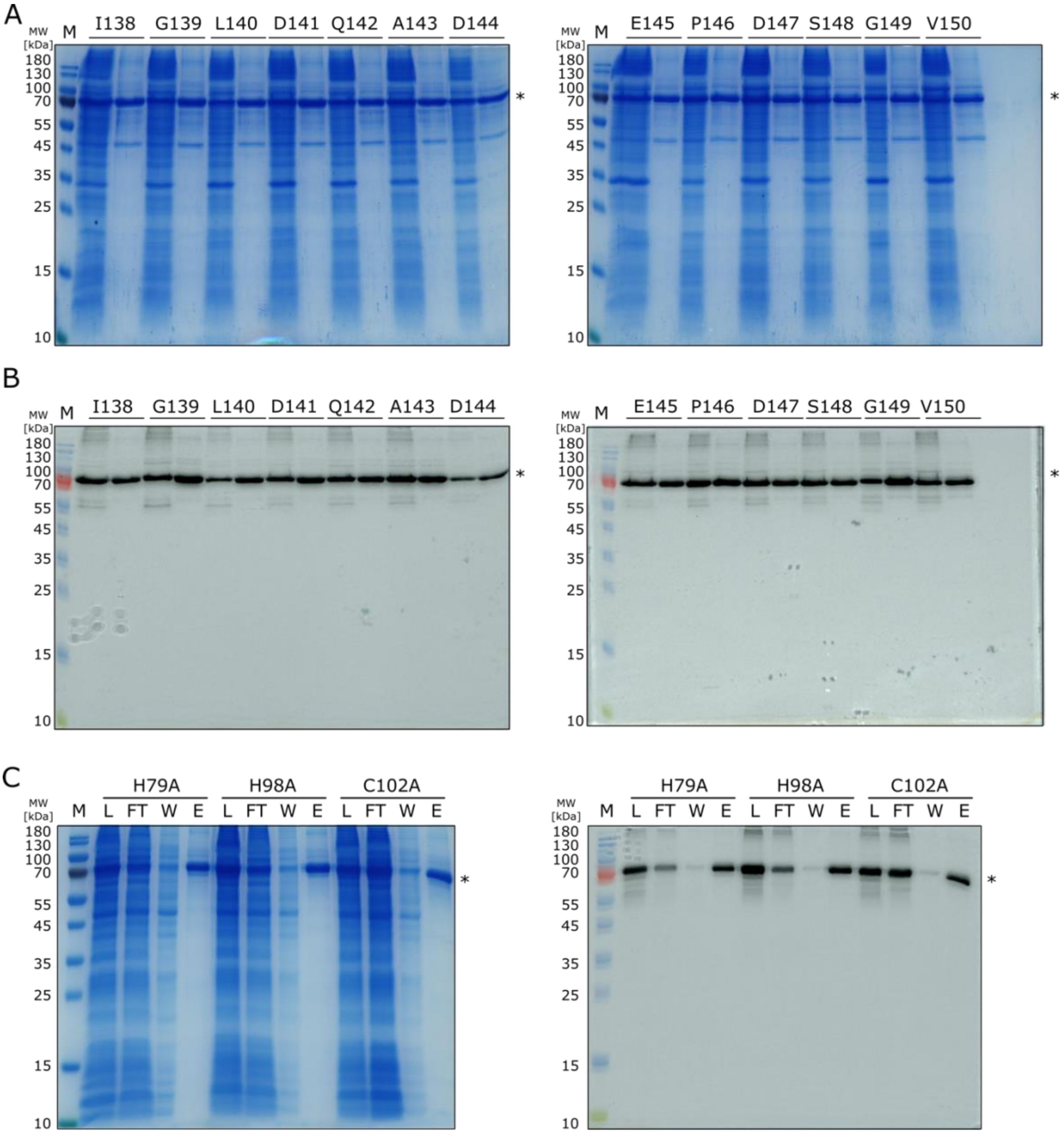


**Figure S2: Expression and Purification of putative biosensor candidates for ferrous iron**

The biosensor expression conducted in bacterial expression host. Cell lysate (left) and the purified biosensors after affinity chromatography analyzed by SDS-PAGE and gel images are for every tested biosensor candidate, (**A**) shows the Coomassie stained images of the SDS-PAGE and (**B**) shows the in-gel fluorescence of the SDS-PAGE of Matryoshka biosensor variants I138-V150, whereby the cell lysate (left) is loaded next to elution sample (right) for each variant. (**C**) Shows the Coomassie stained images and the in-gel fluorescence of the SDS-PAGE of the expression and affinity purification for the binding-deficient mutants of IronSenseR (variant G149) H79A, H98A and C102A. The molecular weight is indicated in kDa. The prominent band at approximately 80 kDa labeled with asterisk (*) indicates the biosensor. L: Cell lysate, FT: Flow through, W: Wash fraction, E: Elution fraction from affinity chromatography.

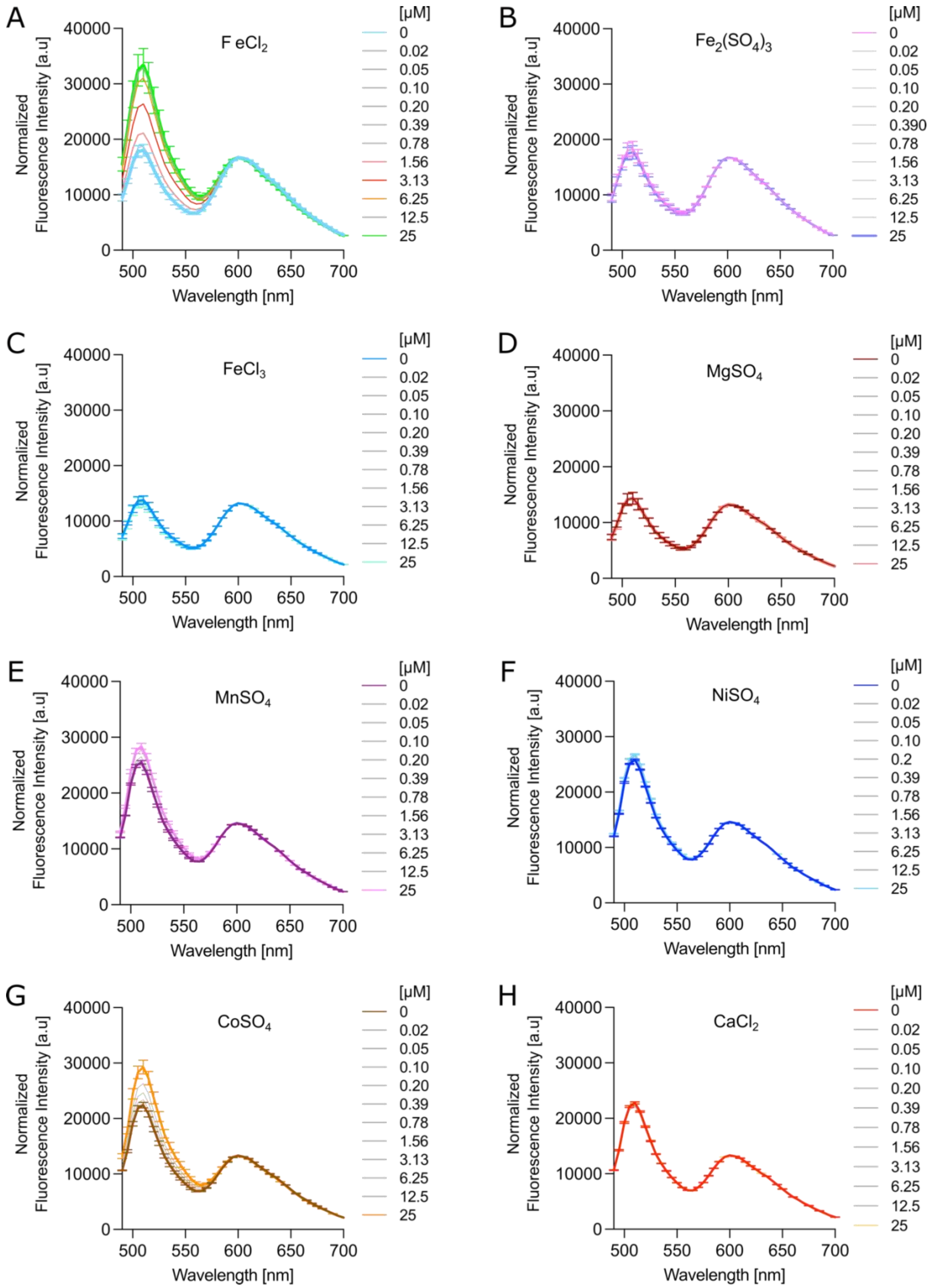


**Figure S3: Fluorescence Spectra of titration of various metal ions to the IronSenseR**

The specificity of IronSenseR for ferrous iron was analyzed *in vitro*. Various metal ions were utilized for titration ranging from 0-25 µM to the biosensor (**A**-**H**). The experiment conducted in biological triplicates (n=3). 0 and 25 µM are colored separately.

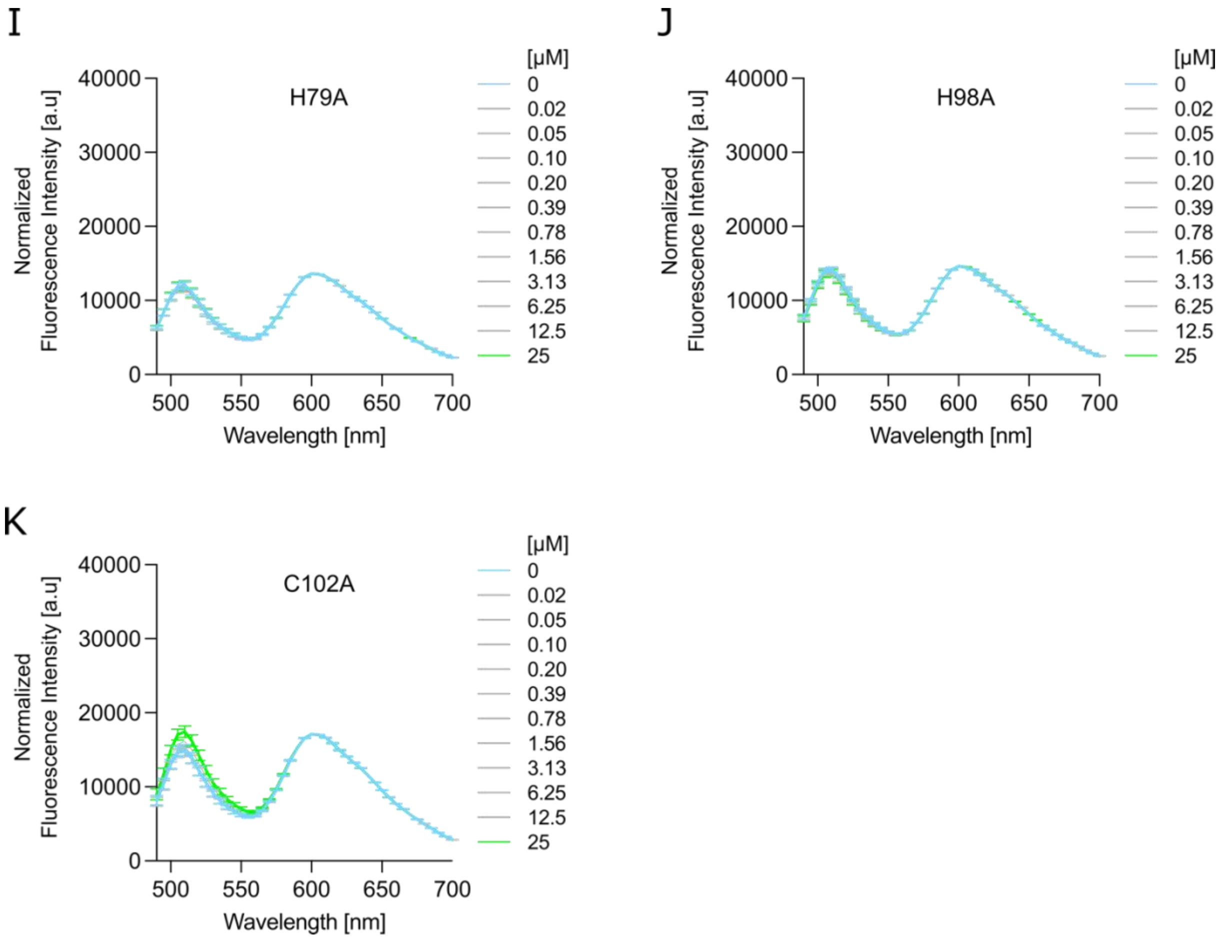


**Figure S3 (continued): Fluorescence Spectra of titration of various metal ions to the IronSenseR**

The sensing action IronSenseR binding deficient mutants H79A, H98A andC102A for ferrous iron was analyzed *in vitro*. Titration ranging from 0-25 µM to the biosensor (**I**-**K**). The experiment conducted in biological triplicates (n=3). 0 and 25 µM are colored separately.

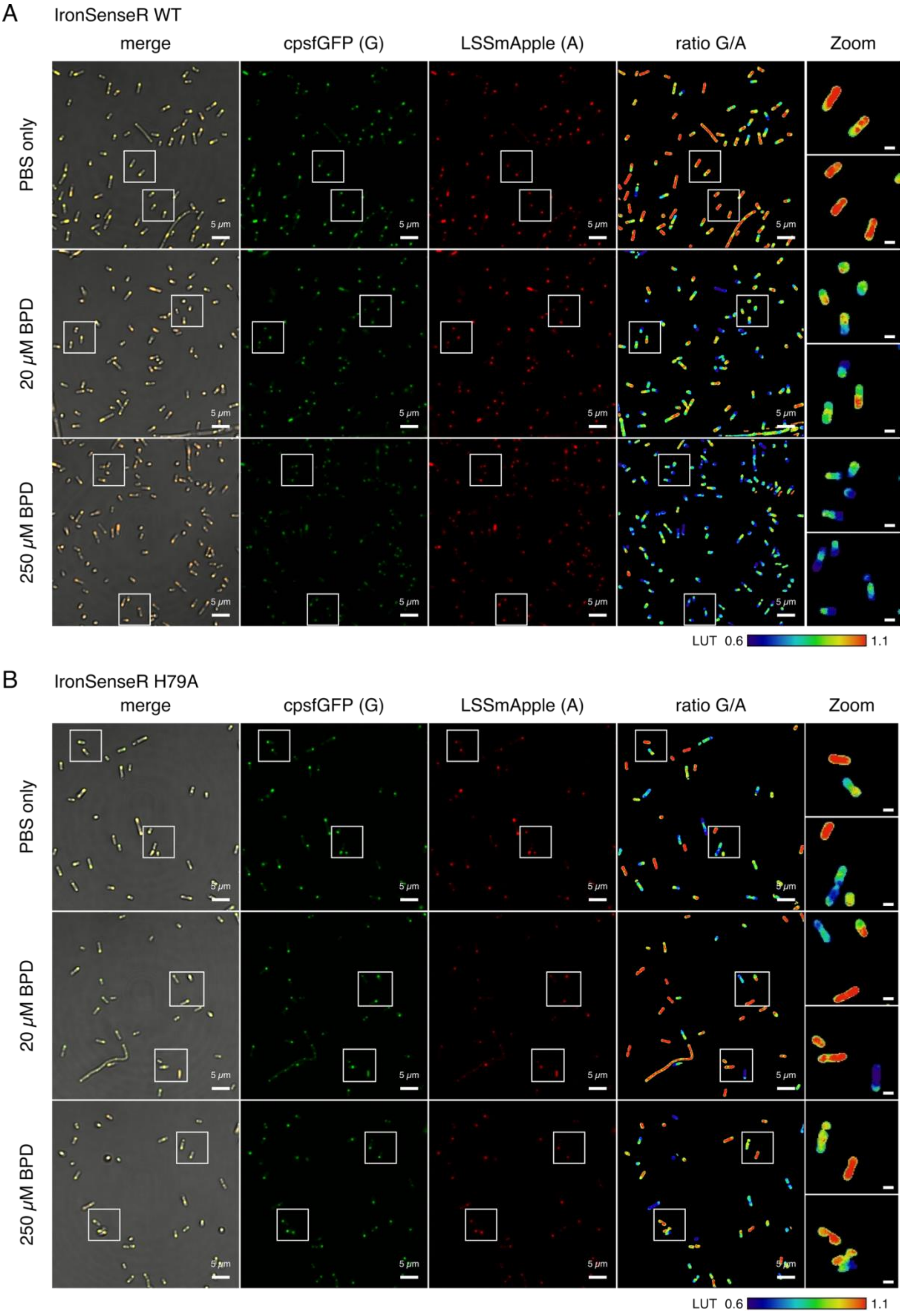


**Figure S4: *In vivo* imaging of *Escherichia coli* expressing IronSenseR variants upon addition of BPD.** Response of IronSenseR WT (A) and binding deficient mutants H79A (B) as well as H98A (C) and C102A (D) to increasing concentrations of the iron chelator 2,2′-Bipyridine (BPD) in bacterial cells. Confocal images of *E. coli* BL21(DE3) expressing the biosensor without addition of BPD (0 µM, PBS top row) and after incubation with either 20 µM BPD (middle row) or 250 µM BPD (bottom row) are depicted. Merged and single fluorescence channels are shown. Samples were excited with a 488 nm laser and cpsfGFP as well as LSSmApple emission were acquired simultaneously at 500-530 nm (cpsfGFP (G)) and 600-700 nm respectively (LSSmApple (A)). Ratio of green (cpsfGFP) to red fluorescence (LSSmApple) is displayed using a rainbow colored lookup table with two zoomed-in sections depicted next to the main column. The scale bar in the zoomed-in images is 1 µm. LUT 0.6-1.1 is depicted by the color bar (down right) indicating the decrease in ratio G/A upon BPD addition. In the case of IronSenseR, when iron is present, ratio G/A and LUT is high (red) but drops (blue) when iron is chelated by BPD. In case of the binding deficient mutant, this effect is not observable/highly diminished. Ratio data indicated also in main figures. Data acquired in biological replicates, n=6 for WT and n=3 for binding deficient mutants.

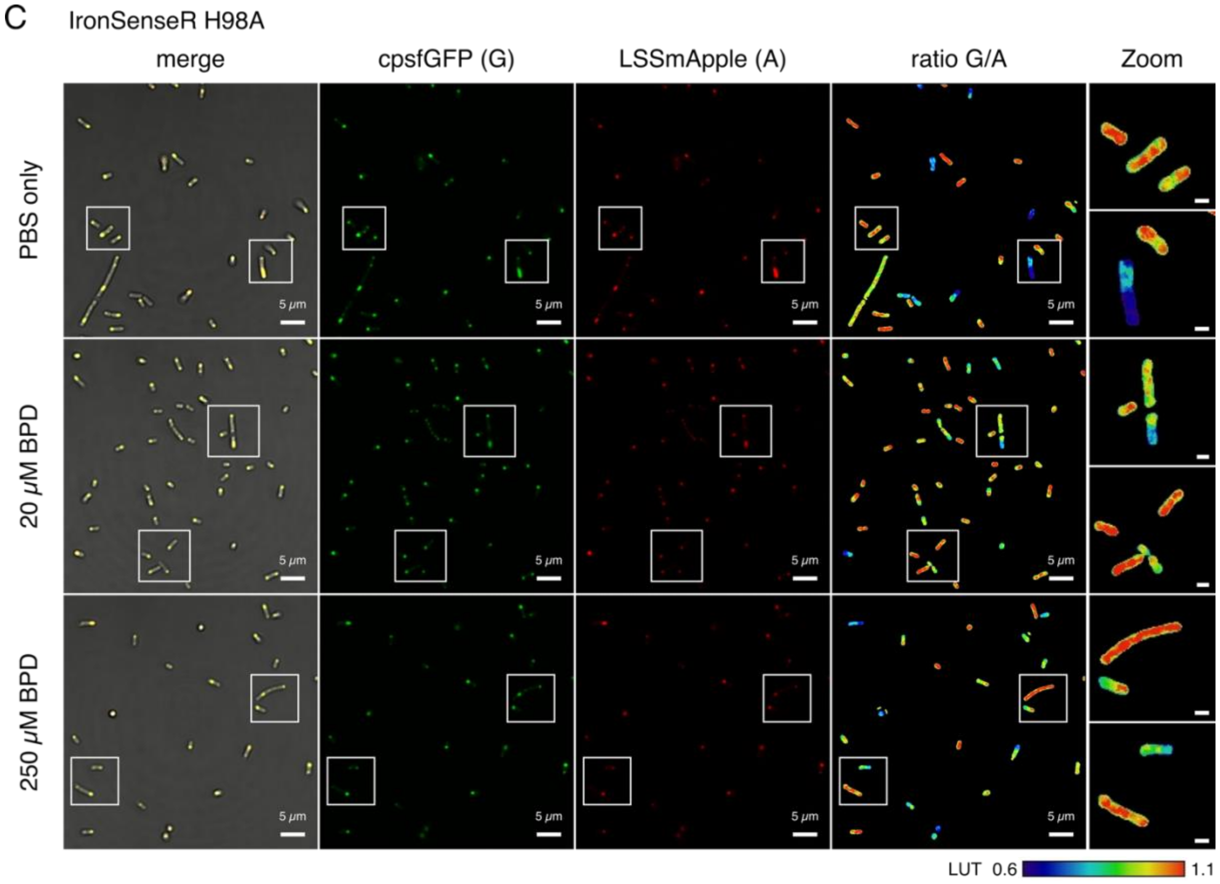


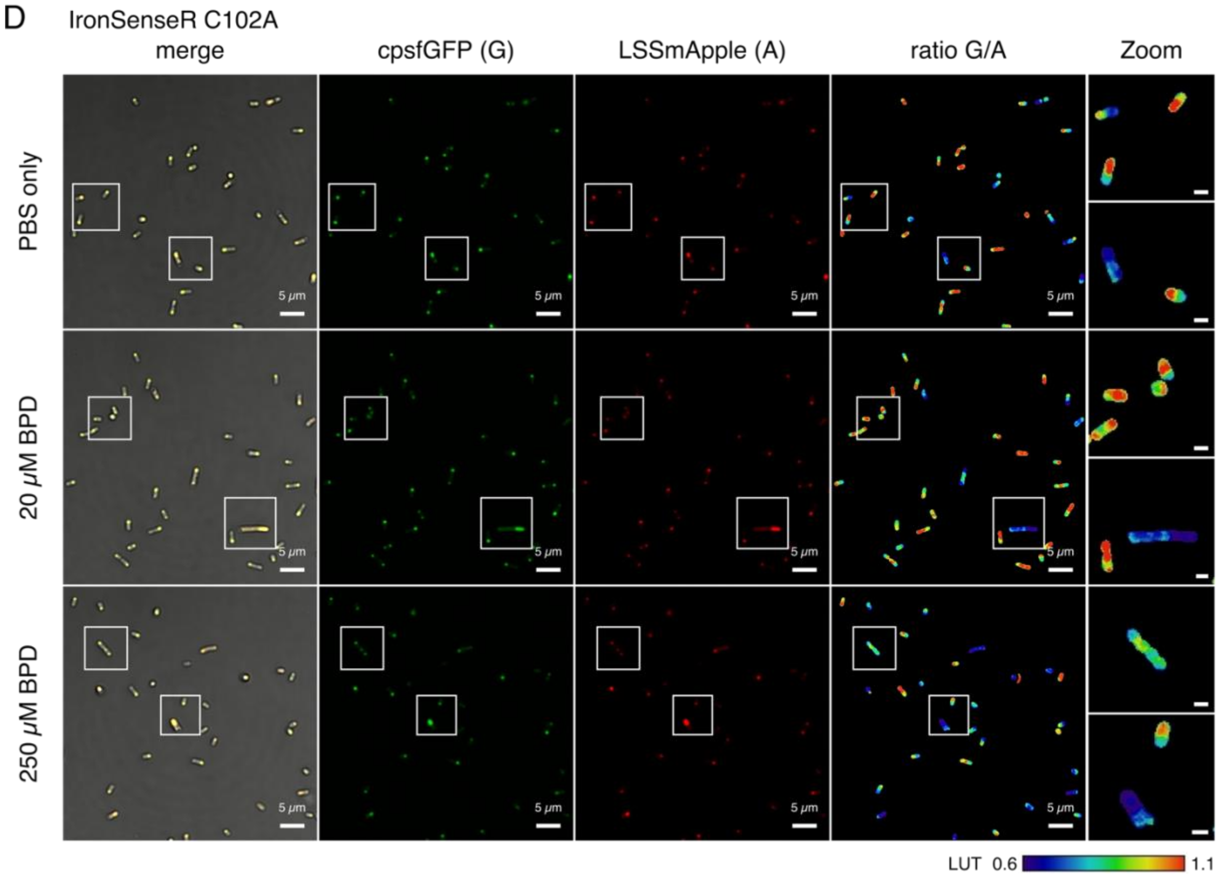


**Figure S4 (continuation):** ***In vivo*** **imaging of** ***Escherichia coli*** **expressing IronSenseR variants upon addition of BPD.**

E

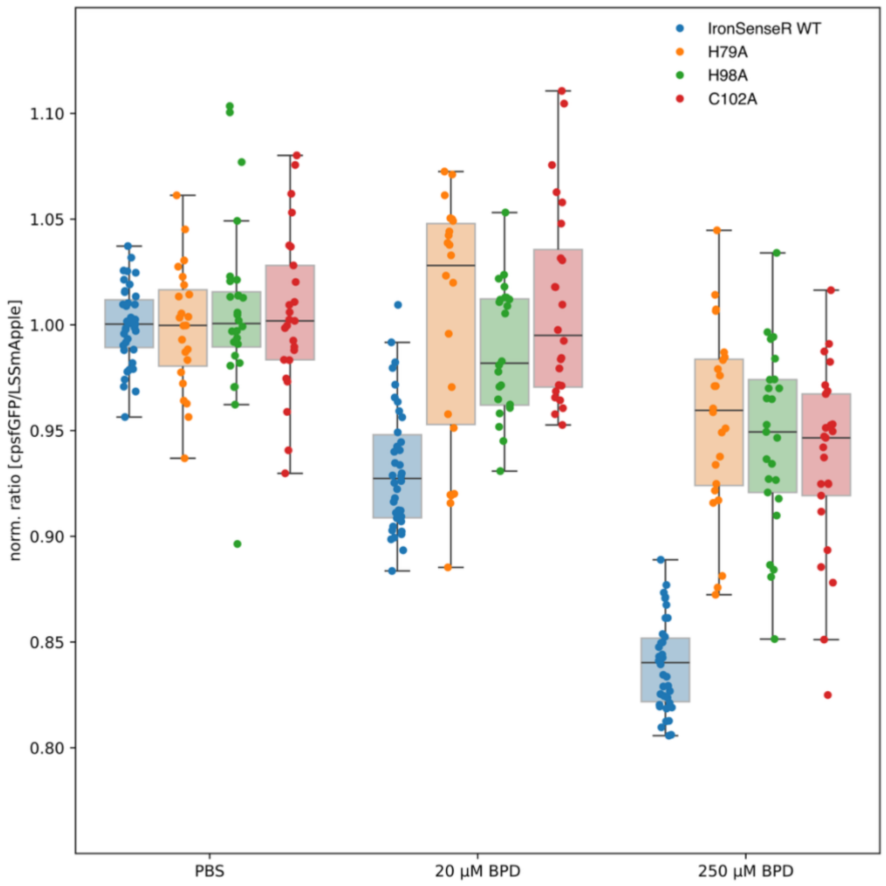


**Figure S4 (continuation):** ***In vivo*** **imaging of** ***Escherichia coli*** **expressing IronSenseR variants upon addition of BPD.** The box plot (E) shows image-wide ratio averages aggregated for different BPD concentration at the time of the reagent addition (24 h). Fluorescence ratios were normalized to the ratio mean of the PBS measurement for each replicate. avg ± std. IronSenseR WT: PBS: 1.00 ± 0.02, 20 µM BPD: 0.93 ± 0.03, 250 µM BPD: 0.84 ± 0.02, H79A: PBS: 1.00 ± 0.03, 20 µM BPD: 1.00 ± 0.06, 250 µM BPD: 0.96 ± 0.05, H98A: PBS: 1.00 ± 0.04, 20 µM BPD: 0.99 ± 0.03, 250 µM BPD: 0.95 ± 0.04, C102A: PBS: 1.00 ± 0.04, 20 µM BPD: 1.01 ± 0.05, 250 µM BPD: 0.94 ± 0.04, IronSenseR WT: n=6, binding deficient mutants (H79A, H98A, C102A): n=3

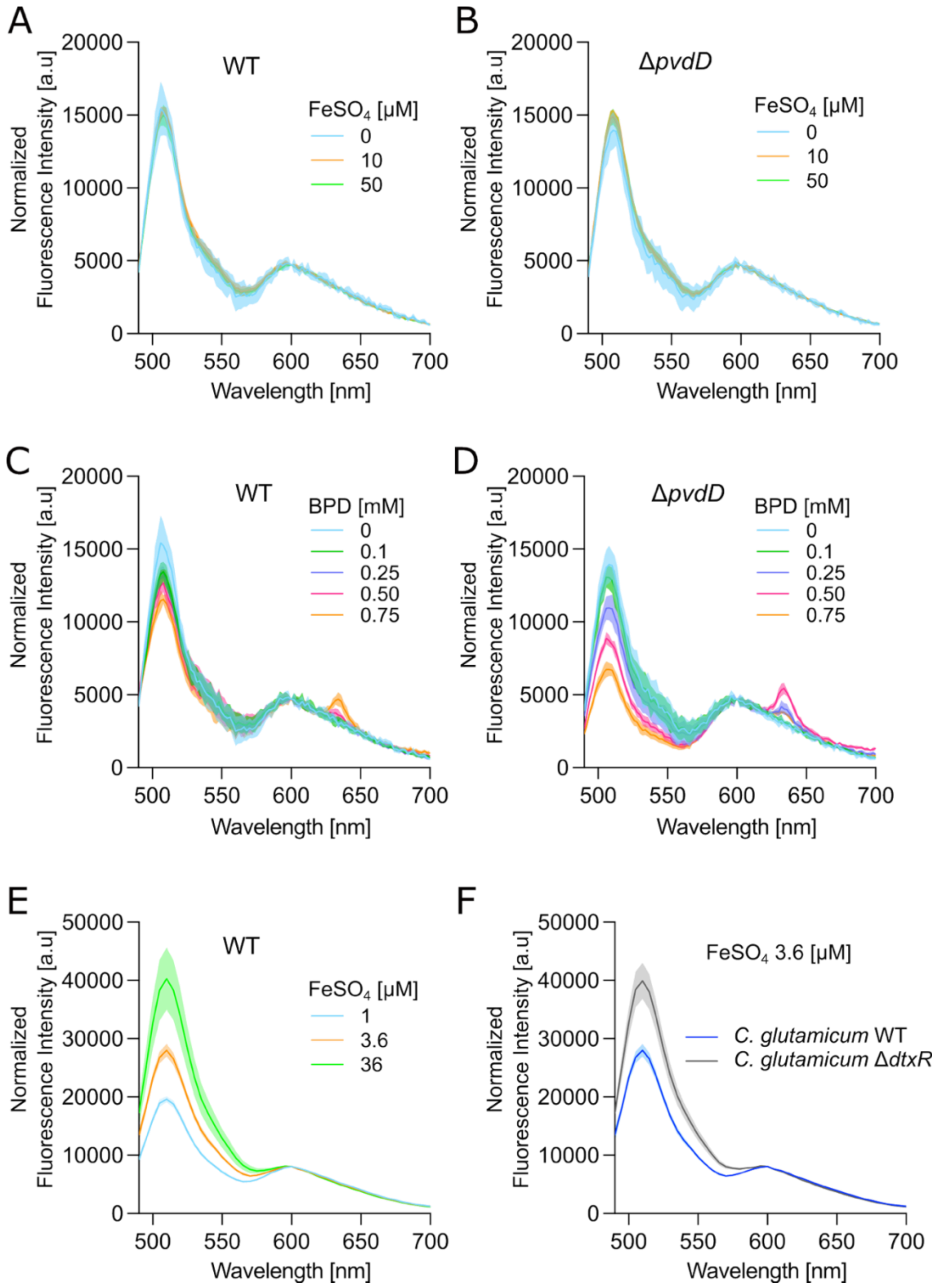


**Figure S5: Spectral analysis of *In vivo* iron sensing in *Pseudomonas putida* and *Corynebacterium glutamicum***

Emission spectra of IronSenseR upon increasing concentrations of the iron chelator 2,2′-Bipyridine (BPD) and ferrous iron ($FeSO_4$) in the *P. putida* and *C. glutamicum*. *P. putida* wild type (WT) and pyoverdine-lacking mutant (Δ*pvdD*) strain as well as *C. glutamicum* WT and DtxR-deletion mutant (*ΔdtxR*) were investigated. **A**: *P. putida* WT cultivated in presence of 0-50 µM ferrous iron ($FeSO_4$) do not indicate changes in IronSenseR fluorescence. **B**: *P. putida* Δ*pvdD* cultivated in presence of 0-50 µM ferrous iron ($FeSO_4$) do not indicate changes in IronSenseR fluorescence. **C**: Cultivation of *P. putida* WT cultivated in presence of 0-0.75 mM BPD did not result in a decrease in the reporter fluorophore signal intensity. These results indicate that the cytoplasmic pool of iron ions remains unchanged under these conditions. **D**: IronSenseR reporter fluorescence decreased with increasing BPD concentration in the mutant strain *P. putida* Δ*pvdD.* In contrast to the WT, this strain is not able to synthesize the iron-chelating siderophore pyoverdine under iron-limiting conditions, resulting in a detectable reduction of the intracellular iron pool. **F**: IronSenseR utilized to sense varying iron content upon cultivation of *C. glutamicum*. Increasing iron uptake and/or availability lead to increasing reporter fluorescence of IronSenseR. **G**: Comparison of *C. glutamicum* WT and Δ*dtxR* strains upon cultivation in 3.6 µM iron and expression of IronSenseR. The mutant strain Δ*dtxR* lacks the regulation of iron acquisition and is not limited in comparison to the WT, therefore IronSenseR signal is increased indicating a large pool of intracellular iron in comparison to the WT. The Experiments conduct in n=6 biological replicates for P. putida and n=4 biological replicates for *C. glutamicum*.

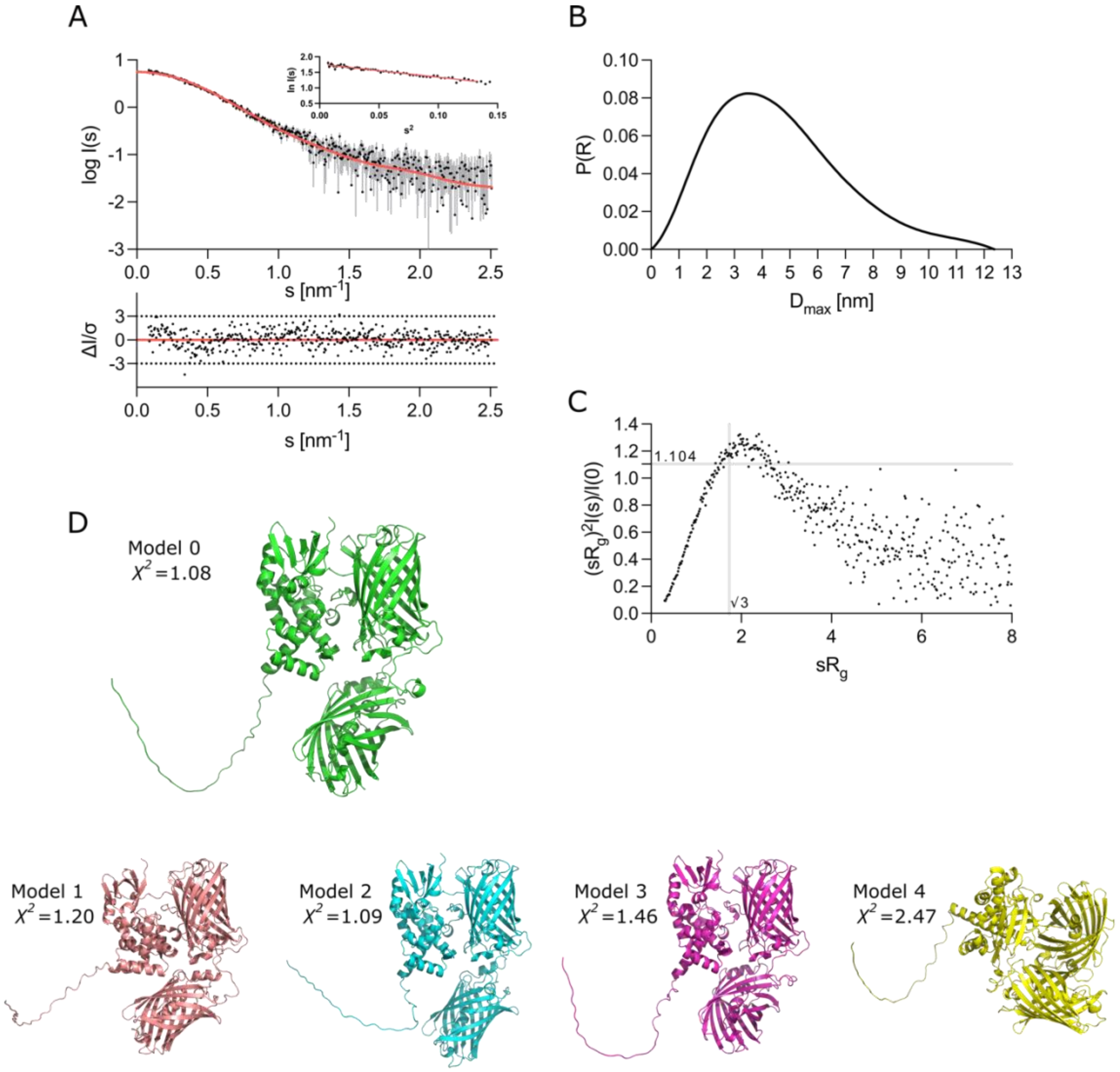


**Figure S5: Small-angle X-ray scattering data from MDtxR$_{G149}$ GA apo. A:** Scattering data of MDtxR$_{G149}$GA apo. Experimental data are shown in black dots, with grey error bars. The theoretical intensity fit, created with CRYSOL is shown as red line and below is the residual plot of the data. The Guinier plot is added in the right corner. **B:** *p(r)* function of MDtxR$_{G149}$GA apo. **C:** Dimensionless Kratky plots of MDtxR$_{G149}$GA apo. **D**: AlphaFold3 models of MDtxR$_{G149}$GA (models 0 – 4) and respective chi square ($X^2$) values against experimental structural data obtained by SAXS.

### Supplemental Tables

**Table S1: Overall SAXS Data**

| | |
|---|---|
| **Data collection parameters** | |
| SAXS Device | BM29, ESRF Grenoble [6] |
| Detector | PILATUS3 x 2 M |
| Detector distance (m) | 2.827 |
| Beam size | 200 µm x 100 µm |
| Wavelength (nm) | 0.099 |
| Sample environment | Quartz glass capillary, 1 mm ø |
| Absolute scaling method | Comparison with scattering from pure $H_2O$ |
| Normalization | To transmitted intensity by beam-stop counter |
| Scattering intensity scale | Absolute scale, $cm^{-1}$ |
| *s* range ($nm^{-1}$), ($s = 4\pi\sin(\theta)/\lambda$) | 0.025–5.5 |
| **Sample** | **IronSenseR apo** |
| Organism | *C. glutamicum* |
| Mode of measurement | SEC-SAXS |
| SEC-Column | Superdex 200 increase 10/300 |
| Flowrate (ml/min) | 0.6 |
| Injection volume (µl) | 100 |
| Temperature (°C) | 20 |
| Exposure time (# frames) | 2 s (1200 frames) |
| # frames used for averaging | 20 |
| Protein buffer | 20 mM MOPS pH 7.0, 250 mM KCl |
| Protein concentration (mg/ml) | 4.00 |
| **Structural parameters** | |
| ***Guinier Analysis (PRIMUS)*** | |
| $I(0) \pm \sigma$ ($cm^{-1}$) | 5.83 ± 0.047 |
| $R_g \pm \sigma$ (nm) | 3.55 ± 0.05 |
| *s-range* ($nm^{-1}$) | 0.084 – 0-364 |
| *min < sRg < max limit* | 0.300 – 1.292 |
| Data point range | 1 - 57 |
| Linear fit assessment ($R^2$) | 0.951 |
| ***PDDF/P(r) Analysis (GNOM 5)*** | |
| $I(0) \pm \sigma$ ($cm^{-1}$) | 5.83 ± 0.042 |
| $R_g \pm \sigma$ (nm) | 3.62 ± 0.03 |
| $D_{max}$ (nm) | 12.37 |
| Porod volume ($nm^3$) | 118.25 |
| *s-range* ($nm^{-1}$) | 0.085 – 2.507 |
| χ2 / CorMap P-value | 0.997 / 0.381 |
| **Molecular mass (kDa)** | |
| From *I*(0) | n.d. |
| From Qp [7] | 74.14 |
| From MoW2 [8] | 85.63 |
| From Vc [9] | 76.87 |
| From Bayesian Inference [10] | 83.13 |
| From Gnnom [11] | 81.40 |
| From sequence | 84.19 (monomer) |
| **Atomistic modelling** | |
| **CRYSOL (with default parameters)** | |
| Constant subtraction allowed | yes |
| *s*-range for fit ($s_{min} – s_{max}$; $nm^{-1}$) | 0.085 – 2.507 |
| $\chi^2$, CorMap *P*-value | 1.086 / 0.384 |
| **SASBDB accession codes [12]** | |
| **Software** | |
| ATSAS Software Version [13] | 3.0.5 |
| Primary data reduction | CHROMIXS [14]/ PRIMUS [15] |
| Data processing | GNOM [16] |
| Atomistic modelling | CRYSOL [17] |
| Model visualization | PyMOL [18] |

‡$s = 4\pi\sin(\theta)/\lambda$, 2θ – scattering angle, λ – Xray-wavelength, n.d. not determined

**Table S2: List of Primers**

| Name | Sequence (5'-3') |
|---|---|
| Construction of $pRSET_B$ 10xHis DtxR and MDtxRGA variants | |
| Pfw_pRSETb_Link_OL | GAAAAGGTTGAGGGCGGCGCCGGTGACGTC |
| Prev_pRSETb_Link_OL | GACCAGATCCTTCATGGCGCCTCCGGTACC |
| Pfw_DtxR_OL | GGTACCGGAGGCGCCATGAAGGATCTGGTC |
| Prev_DtxR_OL | GACGTCACCGGCGCCGCCCTCAACCTTTTC |
| Pfw_Mat_L | CCTGCTAGCCATAACGTGTATATTACC |
| Prev_Mat_L | AGGATTGTTAAAGTTATATTCCAGTTTATGCCC |
| Pfw_DtxR_I138 | TTTAACAATCCTGGTTTGGATCAAGCAGATGAGCCTGAT |
| Prev_DtxR_I138 | ATGGCTAGCAGGGATTTCGCCGAGGCC |
| Pfw_DtxR_G139 | TTTAACAATCCTTTGGATCAAGCAGATGAGCCTGATTCCGGC |
| Prev_DtxR_G139 | ATGGCTAGCAGGACCGATTTCGCCGAG |
| Pfw_DtxR_L140 | TTTAACAATCCTGATCAAGCAGATGAGCCTGATTCCGGC |
| Prev_DtxR_L140 | ATGGCTAGCAGGCAAACCGATTTCGCC |
| Pfw_DtxR_D141 | TTTAACAATCCTCAAGCAGATGAGCCTGATTCCGGCGTT |
| Prev_DtxR_D141 | ATGGCTAGCAGGATCCAAACCGATTTCGCCGAG |
| Pfw_DtxR_Q142 | TTTAACAATCCTGCAGATGAGCCTGATTCCGGCGTT |
| Prev_DtxR_Q142 | ATGGCTAGCAGGTTGATCCAAACCGATTTCGCC |
| Pfw_DtxR_A143 | TTTAACAATCCTGATGAGCCTGATTCCGGCGTTCGTGCC |
| Prev_DtxR_A143 | ATGGCTAGCAGGTGCTTGATCCAAACCGATTTCGCC |
| Pfw_DtxR_D144 | TTTAACAATCCTGAGCCTGATTCCGGCGTTCGT |
| Prev_DtxR_D144 | ATGGCTAGCAGGATCTGCTTGATCCAAACCGATTTC |
| Pfw_DtxR_E145 | TTTAACAATCCTCCTGATTCCGGCGTTCGTGCCATCGATCTG |
| Prev _DtxR_E145 | ATGGCTAGCAGGCTCATCTGCTTGATCCAAACCGAT |
| Pfw_DtxR_P146 | TTTAACAATCCTGATTCCGGCGTTCGTGCCATCGAT |
| Prev_DtxR_P146 | ATGGCTAGCAGGAGGCTCATCTGCTTGATCCAA |
| Pfw_DtxR_D147 | TTTAACAATCCTTCCGGCGTTCGTGCCATCGATCTG |
| Prev_DtxR_D147 | ATGGCTAGCAGGATCAGGCTCATCTGCTTGATCCAA |
| Pfw_DtxR_S148 | TTTAACAATCCTGGCGTTCGTGCCATCGATCTGCCT |
| Prev_DtxR_S148 | ATGGCTAGCAGGGGAATCAGGCTCATCTGCTTG |
| Pfw_DtxR_G149 | TTTAACAATCCTGTTCGTGCCATCGATCTGCCTCTCGGT |
| Prev_DtxR_G149 | ATGGCTAGCAGGGCCGGAATCAGGCTCATC |
| Pfw_DtxR_V150 | TTTAACAATCCTCGTGCCATCGATCTGCCTCTCGGTGAG |
| Prev_DtxR_V150 | ATGGCTAGCAGGAACGCCGGAATCAGGCTC |
| Pfw_DtxR_H79A | GCGCGCCTAGCAGAACGC |
| Prev_DtxR_H79A | CTTACGCATCACGGC |
| Pfw_DtxR_H98A | GCGGACGAAGCATGCCGC |
| Prev_DtxR_H98A | GACTTTGTGGATGTCCAAG |

| | |
|---|---|
| Pfw_DtxR_C102A | CGCTGGGAGCACGTGATGAGT |
| Prev_DtxR_C102A | TGCTGCTTCGTCGTGGACTTTGTG |

Construction of pPREx2-MDtxR$_{G149}$GA

| | |
|---|---|
| MDtxR-G149-GA fw | TGCAGAAGGAGATATACATATGAAGGATCTGGTCGATAC |
| MDtxR-G149-GA rv | AAAACGACGGCCAGTGAATTCCTAGCCCTCAACCTTTTCTACGCG |

Confirmation of integration by colony-PCR

| | |
|---|---|
| Seq1MDtxRG149GA fw | GCCGACATCATAACGGTTCTGG |
| Seq2MDtxRG149GA fw | GAACTTTAAGAAGGAGATATCATATGGTGAGCAAGGGCGAGGAG |
| Seq3MDtxRG149GA rv | CGGCGTTTCACTTCTGAGTTCGGC |

Cloning of mtn7_Ptac_IronSensR in two steps, creating mtn7_Ptac_mCherry in between

| | |
|---|---|
| SP_090_IF_miniTn7_rv | TTAATTAAGACGTCTTGACA |
| SP_091_IF_miniTn7_FW | ATTCGAGCTCGGTACCCGGG |
| SP_092_IF_Ptac_pm_rv | GTACCGAGCTCGAATAAAACTAAAGCGCCACAAGG |
| SP_093_IF_Ptac_pm_fw | GACGTCTTAATTAAAGTCAAAAGCCTCCGGTCGGAG |
| SP_284_IF_GAOPmt_ins_fw | CAGGAAACAGAATTCATGAAAGACCTGGTCGA |
| SP_285_IF_GAOPmt_ins_rv | GAATTTTCTAGAACGCTAGCCTTCGACTTTCTCG |
| SP_286_IF_GAOPmt_BB_fw | AAAGTCGAAGGCTAGCGTTCTAGAAAATTCGTCAA |
| SP_287_IF_GAOPmt_BB_rv | GACCAGGTCTTTCATGAATTCTGTTTCCTGTGTGA |

Cloning of pSNW2_ΔpvdD

| | |
|---|---|
| pvdD-up-fw | CGGAATTCCGTGGCGTATTGCTGGTAG |
| pvdD-up-rv | CGACGCGTCGGCTTTCGGGGCCGCCAGCGCGGCCCCTCTGGAGA ATCGAACG |
| pvdD-dw-fw | CGACGCGTCGTTGAACATCTCCTACCAGGGCACCGGTCTTTG |
| pvdD-dw-rv | GCTCTAGAGCTATCTGCGTGCCAGCCTTC |

**Table S3: Bacterial strains utilized in this study**

| Strain | Relevant characteristics |
|---|---|
| ***Escherichia coli*** | |
| DH5α[19] | F⁻ ϕ80*dlac* Δ(*lacZ*)M15 Δ(*lacZYA-argF*) U169 *endA1 recA1 hsdR17* ($r_K^-$ $m_K^+$) *deoR thi-1 phoA supE44* λ⁻ *gyrA96 relA1*; strain used for cloning procedures |
| BL21 (DE3)[20] | *E. coli* str. B F– *ompT gal dcm lon hsdSB* (*rB–mB–*) λ (DE3) [*lacI lacUV5-T7p07 ind1 sam7 nin5*]) [*malB*+]K-12 (λS); used for biosensor expresion |
| HB101[21] | *F− mcrB mrr hsdS20(rB− mB−) recA13 leuB6 ara-14 proA2 lacY1 galK2 xyl-5 mtl-1 rpsL20(SmR) gln V44λ−* |
| PIR2 (Life techonologies) | *F- Δlac169 rpoS (Am) robA1 creC510 hsdR514 endA reacA1 uidA (ΔMlui)::pir* |
| DH5α λpir[22] | *endA1 hsdR17 glnV44 (= supE44) thi-1 recA1 gyrA96 relA1 φ80dlacΔ(lacZ)M15 Δ(lacZYA-argF)U169 zdg-232::Tn10 uidA::pir*+ |
| ***Pseudomonas putida*** | |
| KT2440[23] | wild type, derivative of mt-2 |
| Δ*pvdD* | *P. putida* KT2440 with deletion of *pvdD* gene |
| IronSensR | *P. putida* KT2440 with insertion of the IronSensR cassette into the attTn7-site |
| Δ*pvdD*-IronSensR | *P. putida* Δ*pvdD* with insertion of the IronSensR cassette into the attTn7-site |
| ***Corynebacterium glutamicum*** | |
| ATCC13032 (WT)[24] | Wild type, biotin auxotrophic |
| ATCC13032 Δ*dtxR*[25] | In-frame deletion of the *dtxR* gene |

**Table S4: List of plasmids**

| Strain | Relevant characteristics |
|---|---|
| pAZ180-191 | Amp, $pRSET_B$ derivative for expression of the matryoshka cassette (cpsfGFP and nested LSSmApple inserted into the respective insertion sites of DtxR (I138-V150)) with an N-terminal 10x-His-tag, T7-tag and Xpress-tag under the control of T7 promoter, Enterokinase cleavage site |
| pPREx2[26] | $Kan^r$, pPBEx2 derivative ($P_{tacI}$, $lacI^q$, $ori_{C.g}$ from pBL1.; $ori_{E.c.}$ ColE1 from pUC18), with a consensus RBS (AAGGAG) for *C. glutamicum* |
| pPREx2-$MDtxR_{G149}$GA | $Kan^r$, pPREx2 derivative carrying the Matryoshka cassette, encoding for the cpsfGFP and nested LSSmApple inserted into the respective insertion sites of DtxR (I138-V150) under the control of $P_{tac}$ promoter |
| pRK2013[27] | $Km^R$, *oriV*(RK2/ColE1) -*mob*$^+$ *tra*$^+$ |
| pTNS1[28] | $Ap^R$, *oriV*(R6K), *TnSABC+D* operon |
| pBG13[29] | $Km^R$, $Gm^R$, *ori*R6K, pBG-derived, promoter $P_{em7}$, *msfGFP* |
| mtn7_Ptac_mCherry | $Km^R$, $Gm^R$, *ori*R6K, $P_{tac}$, *lacI, mCherry* |
| mtn7_Ptac_IronSensR | $Km^R$, $Gm^R$, *ori*R6K, $P_{tac}$, *lacI,* IronSensR |
| pSNW2[30] | Suicide vector used for deletions in Gram-negative bacteria; *oriT, traJ, lacZα, ori*(R6K), $P_{14g}$(*BCD2*)→*msfGFP*; $Km^R$ |
| pSNW2_ΔpvdD | pSNW2 , bearing flanking sequences of *pvdD* |
| pQURE6·H[30] | *oriV*(RK2), XylS/Pm→*I-SceI* and $P_{14g}$*(BCD2)*→*mRFP*; $Gm^R$ |

### Supplemental Sequences

Codon optimized sequences for IronSenseR in Pseudomonas putida

>IronSenseR CO Pp
ATGAAAGACCTGGTCGACACCACCGAGATGTACCTGCGCACGATCTACGAACTGGAGG
AGGAAGGCATCGTCCCACTGCGCGCGCGCATCGCGGAGCGCCTCGAACAGTCGGGCC
CAACGGTCAGCCAGACGGTCGCCCGCATGGAGCGCGATGGCCTCGTGCACGTCAGCC
CTGATCGCAGCTTGGAGATGACGCCAGAAGGGCGCTCGTTGGCCATCGCCGTGATGC
GCAAACACCGCCTTGCGGAGCGCTTGCTCACCGATATCATCGGCTTGGATATCCACAA
GGTCCACGATGAGGCGTGCCGCTGGGAACACGTGATGTCGGACGAAGTCGAGCGCCG
CTTGGTCGAGGTCCTCGATGACGTGCACCGCTCGCCATTCGGCAACCCAATCCCAGGC
TTGGGCGAGATCGGCTTGGACCAAGCGGACGAACCAGACTCGGGCCCAGCGAGCCAC
AACGTGTACATCACCGCCGACAAGCAGAAGAACGGCATCAAGGCCAACTTCACCGTGC
GCCACAACGTGGAGGACGGCAGCGTGCAGCTGGCCGACCACTACCAGCAGAACACCC
CGATCGGCGACGGCCCGGTGCTGCTGCCGGACAACCACTACCTGAGCACCCAGACCA
AACTGAGCAAGGACCCGAACGAGAAGCGCGACCACATGGTGCTGCTGGAGTTCGTGAC
CGCGGCCGGCATCACGCACGGCATGGACGAGCTGTACGGCGGCACCGTGAGCAAAGG
CGAAGAAAACAACATGGCCATCATCAAAGAATTCATGCGCTTCAAAGTGCACATGGAAG
GCTCGGTGAACGGCCACGAATTCGAAATCGAAGGCGAAGGCGAAGGCCGCCCTTACG
AAGCCTTCCAGACCGCGAAACTGAAAGTGACCAAAGGCGGCCCTCTGCCTTTCACCTG
GGATATCCTGTCGCCACAGTTCATGTACGGCTCGAAAGTCTACATCAAACACCCAGCCG
ATATCCCTGATTACTTCAAACTGTCGTTCCCTGAAGGCTTCAGGTGGGAACGCGTGATG
ATCTTCGAAGATGGCGGCATCATCCACGTCAACCAGGATTCGTCGCTGCAGGATGGCG
TGTTCATCTACAAAGTGAAACTGCGCGGCACCAACTTCCCTTCGGATGGCCCTGTCATG
CAGAAAAAAACCATGGGCCTCGAAGCCTGCGAAGAACGGATGTACCCTGAAGATGGCG
CCCTGAAAAGCGAATACAAAGAATGGCTGAAACTGAAAGATGGCGGCCACTACGCCGC
CGAAGTCAAAACCACCTACAAAGCCAAAAAACCTGTGCAGCTGCCAGGCGCCTACATC
GTCGATATCAAATTGGATATCGTGTCGCACAACGAAGATTACACCATCGTGGAGCAGTA
CGAGCGCGCCGAAGGCCGCCACTCGACCGGCGGCATGGATGAACTGTACAAAGGCGG
CAGCGCCAGCCAGGGCGAGGAGCTGTTCACCGGCGTGGTGCCGATCCTGGTGGAGCT
GGACGGCGACGTGAACGGCCACAAGTTCAGCGTGCGCGGCGAGGGCGAGGGCGACG
CCACCATCGGCAAGCTGACCCTGAAGTTCATCTCGACCACCGGCAAGCTTCCGGTGCC
GTGGCCGACCCTGGTGACCACCTTGACCTACGGCGTGCAGTGCTTCAGCCGCTACCCG
GACCACATGAAGCGCCACGACTTCTTCAAGAGCGCCATGCCGGAGGGCTACGTGCAGG
AGCGCACCATCAGCTTCAAGGACGACGGCAAGTACAAGACCCGCGCCGTGGTGAAGTT
CGAGGGCGACACCCTGGTGAACCGCATCGAGCTGAAGGGCACCGACTTCAAGGAGGA
CGGCAACATCCTGGGTCACAAGCTGGAGTACAACTTCAACAACCCAGTCCGCGCCATC
GACCTGCCATTGGGCGAAAACCTGAAAGCGCGCATCGTCCAGTTGAACGAAATCCTGC
AGGTCGACCTCGAACAGTTCCAGGCGTTGACCGATGCCGGCGTCGAGATCGGCACCG
AGGTCGATATCATCAACGAACAGGGCCGGGTCGTGATCACCCACAACGGCTCGAGCGT
CGAGCTGATCGATGACCTCGCGCACGCGGTCCGCGTCGAGAAAGTCGAAGGCTAG

The FASTA sequences are depicted below:

Metabolite binding protein
>sp|Q8NP95|DTXR_CORGL Diphtheria toxin repressor OS=Corynebacterium glutamicum (strain ATCC 13032 / DSM 20300 / JCM 1318 / BCRC 11384 / CCUG 27702 / LMG 3730 / NBRC 12168 / NCIMB 10025 / NRRL B-2784 / 534) OX=196627 GN=dtxR PE=3 SV=1
MKDLVDTTEMYLRTIYELEEEGIVPLRARIAERLEQSGPTVSQTVARMERDGLVHVSPDRSL
EMTPEGRSLAIAVMRKHRLAERLLTDIIGLDIHKVHDEACRWEHVMSDEVERRLVEVLDDVH
RSPFGNPIPGLGEIGLDQADEPDSGVRAIDLPLGENLKARIVQLNEILQVDLEQFQALTDAGV
EIGTEVDIINEQGRVVITHNGSSVELIDDLAHAVRVEKVEG

Biosensor Variants
>pAZ180_pRSET_10xHis_EK_MDtxRGA_I138 (749 aa)
MRGSHHHHHHHHHHGMASMTGGQQMGRDLYDDDDKDPGRMKDLVDTTEMYLRTIYELE
EEGIVPLRARIAERLEQSGPTVSQTVARMERDGLVHVSPDRSLEMTPEGRSLAIAVMRKHRL
AERLLTDIIGLDIHKVHDEACRWEHVMSDEVERRLVEVLDDVHRSPFGNPIPGLGEIPASHNV
YITADKQKNGIKANFTVRHNVEDGSVQLADHYQQNTPIGDGPVLLPDNHYLSTQTKLSKDPN
EKRDHMVLLEFVTAAGITHGMDELYGGTVSKGEENNMAIIKEFMRFKVHMEGSVNGHEFEI
EGEGEGRPYEAFQTAKLKVTKGGPLPFTWDILSPQFMYGSKVYIKHPADIPDYFKLSFPEGF
RWERVMIFEDGGIIHVNQDSSLQDGVFIYKVKLRGTNFPSDGPVMQKKTMGLEACEERMYP
EDGALKSEYKEWLKLKDGGHYAAEVKTTYKAKKPVQLPGAYIVDIKLDIVSHNEDYTIVEQYE
RAEGRHSTGGMDELYKGGSASQGEELFTGVVPILVELDGDVNGHKFSVRGEGEGDATIGK
LTLKFISTTGKLPVPWPTLVTTLTYGVQCFSRYPDHMKRHDFFKSAMPEGYVQERTISFKDD
GKYKTRAVVKFEGDTLVNRIELKGTDFKEDGNILGHKLEYNFNNPGLDQADEPDSGVRAIDL
PLGENLKARIVQLNEILQVDLEQFQALTDAGVEIGTEVDIINEQGRVVITHNGSSVELIDDLAH
AVRVEKVEG*

>pAZ181_pRSET_10xHis_EK_MDtxRGA_G139 (749 aa)
MRGSHHHHHHHHHHGMASMTGGQQMGRDLYDDDDKDPGRMKDLVDTTEMYLRTIYELE
EEGIVPLRARIAERLEQSGPTVSQTVARMERDGLVHVSPDRSLEMTPEGRSLAIAVMRKHRL
AERLLTDIIGLDIHKVHDEACRWEHVMSDEVERRLVEVLDDVHRSPFGNPIPGLGEIGPASH
NVYITADKQKNGIKANFTVRHNVEDGSVQLADHYQQNTPIGDGPVLLPDNHYLSTQTKLSKD
PNEKRDHMVLLEFVTAAGITHGMDELYGGTVSKGEENNMAIIKEFMRFKVHMEGSVNGHEF
EIEGEGEGRPYEAFQTAKLKVTKGGPLPFTWDILSPQFMYGSKVYIKHPADIPDYFKLSFPE
GFRWERVMIFEDGGIIHVNQDSSLQDGVFIYKVKLRGTNFPSDGPVMQKKTMGLEACEERM
YPEDGALKSEYKEWLKLKDGGHYAAEVKTTYKAKKPVQLPGAYIVDIKLDIVSHNEDYTIVEQ
YERAEGRHSTGGMDELYKGGSASQGEELFTGVVPILVELDGDVNGHKFSVRGEGEGDATI
GKLTLKFISTTGKLPVPWPTLVTTLTYGVQCFSRYPDHMKRHDFFKSAMPEGYVQERTISFK
DDGKYKTRAVVKFEGDTLVNRIELKGTDFKEDGNILGHKLEYNFNNPLDQADEPDSGVRAID
LPLGENLKARIVQLNEILQVDLEQFQALTDAGVEIGTEVDIINEQGRVVITHNGSSVELIDDLAH
AVRVEKVEG*

>pAZ182_pRSET_10xHis_EK_MDtxRGA_L140 (749 aa)
MRGSHHHHHHHHHHGMASMTGGQQMGRDLYDDDDKDPGRMKDLVDTTEMYLRTIYELE
EEGIVPLRARIAERLEQSGPTVSQTVARMERDGLVHVSPDRSLEMTPEGRSLAIAVMRKHRL
AERLLTDIIGLDIHKVHDEACRWEHVMSDEVERRLVEVLDDVHRSPFGNPIPGLGEIGLPASH
NVYITADKQKNGIKANFTVRHNVEDGSVQLADHYQQNTPIGDGPVLLPDNHYLSTQTKLSKD
PNEKRDHMVLLEFVTAAGITHGMDELYGGTVSKGEENNMAIIKEFMRFKVHMEGSVNGHEF
EIEGEGEGRPYEAFQTAKLKVTKGGPLPFTWDILSPQFMYGSKVYIKHPADIPDYFKLSFPE

GFRWERVMIFEDGGIIHVNQDSSLQDGVFIYKVKLRGTNFPSDGPVMQKKTMGLEACEERM
YPEDGALKSEYKEWLKLKDGGHYAAEVKTTYKAKKPVQLPGAYIVDIKLDIVSHNEDYTIVEQ
YERAEGRHSTGGMDELYKGGSASQGEELFTGVVPILVELDGDVNGHKFSVRGEGEGDATI
GKLTLKFISTTGKLPVPWPTLVTTLTYGVQCFSRYPDHMKRHDFFKSAMPEGYVQERTISFK
DDGKYKTRAVVKFEGDTLVNRIELKGTDFKEDGNILGHKLEYNFNNPDQADEPDSGVRAIDL
PLGENLKARIVQLNEILQVDLEQFQALTDAGVEIGTEVDIINEQGRVVITHNGSSVELIDDLAH
AVRVEKVEG*

>pAZ183_pRSET_10xHis_EK_MDtxRGA_D141  (749 aa)
MRGSHHHHHHHHHHGMASMTGGQQMGRDLYDDDDKDPGRMKDLVDTTEMYLRTIYELE
EEGIVPLRARIAERLEQSGPTVSQTVARMERDGLVHVSPDRSLEMTPEGRSLAIAVMRKHRL
AERLLTDIIGLDIHKVHDEACRWEHVMSDEVERRLVEVLDDVHRSPFGNPIPGLGEIGLDPAS
HNVYITADKQKNGIKANFTVRHNVEDGSVQLADHYQQNTPIGDGPVLLPDNHYLSTQTKLSK
DPNEKRDHMVLLEFVTAAGITHGMDELYGGTVSKGEENNMAIIKEFMRFKVHMEGSVNGHE
FEIEGEGEGRPYEAFQTAKLKVTKGGPLPFTWDILSPQFMYGSKVYIKHPADIPDYFKLSFPE
GFRWERVMIFEDGGIIHVNQDSSLQDGVFIYKVKLRGTNFPSDGPVMQKKTMGLEACEERM
YPEDGALKSEYKEWLKLKDGGHYAAEVKTTYKAKKPVQLPGAYIVDIKLDIVSHNEDYTIVEQ
YERAEGRHSTGGMDELYKGGSASQGEELFTGVVPILVELDGDVNGHKFSVRGEGEGDATI
GKLTLKFISTTGKLPVPWPTLVTTLTYGVQCFSRYPDHMKRHDFFKSAMPEGYVQERTISFK
DDGKYKTRAVVKFEGDTLVNRIELKGTDFKEDGNILGHKLEYNFNNPQADEPDSGVRAIDLP
LGENLKARIVQLNEILQVDLEQFQALTDAGVEIGTEVDIINEQGRVVITHNGSSVELIDDLAHA
VRVEKVEG*

>pAZ184_pRSET_10xHis_EK_MDtxRGA_Q142  (749 aa)
MRGSHHHHHHHHHHGMASMTGGQQMGRDLYDDDDKDPGRMKDLVDTTEMYLRTIYELE
EEGIVPLRARIAERLEQSGPTVSQTVARMERDGLVHVSPDRSLEMTPEGRSLAIAVMRKHRL
AERLLTDIIGLDIHKVHDEACRWEHVMSDEVERRLVEVLDDVHRSPFGNPIPGLGEIGLDQP
ASHNVYITADKQKNGIKANFTVRHNVEDGSVQLADHYQQNTPIGDGPVLLPDNHYLSTQTKL
SKDPNEKRDHMVLLEFVTAAGITHGMDELYGGTVSKGEENNMAIIKEFMRFKVHMEGSVNG
HEFEIEGEGEGRPYEAFQTAKLKVTKGGPLPFTWDILSPQFMYGSKVYIKHPADIPDYFKLSF
PEGFRWERVMIFEDGGIIHVNQDSSLQDGVFIYKVKLRGTNFPSDGPVMQKKTMGLEACEE
RMYPEDGALKSEYKEWLKLKDGGHYAAEVKTTYKAKKPVQLPGAYIVDIKLDIVSHNEDYTIV
EQYERAEGRHSTGGMDELYKGGSASQGEELFTGVVPILVELDGDVNGHKFSVRGEGEGDA
TIGKLTLKFISTTGKLPVPWPTLVTTLTYGVQCFSRYPDHMKRHDFFKSAMPEGYVQERTISF
KDDGKYKTRAVVKFEGDTLVNRIELKGTDFKEDGNILGHKLEYNFNNPADEPDSGVRAIDLP
LGENLKARIVQLNEILQVDLEQFQALTDAGVEIGTEVDIINEQGRVVITHNGSSVELIDDLAHA
VRVEKVEG*

>pAZ185_pRSET_10xHis_EK_MDtxRGA_A143  (749 aa)
MRGSHHHHHHHHHHGMASMTGGQQMGRDLYDDDDKDPGRMKDLVDTTEMYLRTIYELE
EEGIVPLRARIAERLEQSGPTVSQTVARMERDGLVHVSPDRSLEMTPEGRSLAIAVMRKHRL
AERLLTDIIGLDIHKVHDEACRWEHVMSDEVERRLVEVLDDVHRSPFGNPIPGLGEIGLDQA
PASHNVYITADKQKNGIKANFTVRHNVEDGSVQLADHYQQNTPIGDGPVLLPDNHYLSTQT
KLSKDPNEKRDHMVLLEFVTAAGITHGMDELYGGTVSKGEENNMAIIKEFMRFKVHMEGSV
NGHEFEIEGEGEGRPYEAFQTAKLKVTKGGPLPFTWDILSPQFMYGSKVYIKHPADIPDYFK
LSFPEGFRWERVMIFEDGGIIHVNQDSSLQDGVFIYKVKLRGTNFPSDGPVMQKKTMGLEA
CEERMYPEDGALKSEYKEWLKLKDGGHYAAEVKTTYKAKKPVQLPGAYIVDIKLDIVSHNED
YTIVEQYERAEGRHSTGGMDELYKGGSASQGEELFTGVVPILVELDGDVNGHKFSVRGEG
EGDATIGKLTLKFISTTGKLPVPWPTLVTTLTYGVQCFSRYPDHMKRHDFFKSAMPEGYVQE

RTISFKDDGKYKTRAVVKFEGDTLVNRIELKGTDFKEDGNILGHKLEYNFNNPDEPDSGVRAI
DLPLGENLKARIVQLNEILQVDLEQFQALTDAGVEIGTEVDIINEQGRVVITHNGSSVELIDDLA
HAVRVEKVEG*

>pAZ186_pRSET_10xHis_EK_MDtxRGA_D144 (749 aa)
MRGSHHHHHHHHHHGMASMTGGQQMGRDLYDDDDKDPGRMKDLVDTTEMYLRTIYELE
EEGIVPLRARIAERLEQSGPTVSQTVARMERDGLVHVSPDRSLEMTPEGRSLAIAVMRKHRL
AERLLTDIIGLDIHKVHDEACRWEHVMSDEVERRLVEVLDDVHRSPFGNPIPGLGEIGLDQA
DPASHNVYITADKQKNGIKANFTVRHNVEDGSVQLADHYQQNTPIGDGPVLLPDNHYLSTQ
TKLSKDPNEKRDHMVLLEFVTAAGITHGMDELYGGTVSKGEENNMAIIKEFMRFKVHMEGS
VNGHEFEIEGEGEGRPYEAFQTAKLKVTKGGPLPFTWDILSPQFMYGSKVYIKHPADIPDYF
KLSFPEGFRWERVMIFEDGGIIHVNQDSSLQDGVFIYKVKLRGTNFPSDGPVMQKKTMGLE
ACEERMYPEDGALKSEYKEWLKLKDGGHYAAEVKTTYKAKKPVQLPGAYIVDIKLDIVSHNE
DYTIVEQYERAEGRHSTGGMDELYKGGSASQGEELFTGVVPILVELDGDVNGHKFSVRGE
GEGDATIGKLTLKFISTTGKLPVPWPTLVTTLTYGVQCFSRYPDHMKRHDFFKSAMPEGYV
QERTISFKDDGKYKTRAVVKFEGDTLVNRIELKGTDFKEDGNILGHKLEYNFNNPEPDSGVR
AIDLPLGENLKARIVQLNEILQVDLEQFQALTDAGVEIGTEVDIINEQGRVVITHNGSSVELIDD
LAHAVRVEKVEG*

>pAZ187_pRSET_10xHis_EK_MDtxRGA_E145 (749 aa)
MRGSHHHHHHHHHHGMASMTGGQQMGRDLYDDDDKDPGRMKDLVDTTEMYLRTIYELE
EEGIVPLRARIAERLEQSGPTVSQTVARMERDGLVHVSPDRSLEMTPEGRSLAIAVMRKHRL
AERLLTDIIGLDIHKVHDEACRWEHVMSDEVERRLVEVLDDVHRSPFGNPIPGLGEIGLDQA
DEPASHNVYITADKQKNGIKANFTVRHNVEDGSVQLADHYQQNTPIGDGPVLLPDNHYLST
QTKLSKDPNEKRDHMVLLEFVTAAGITHGMDELYGGTVSKGEENNMAIIKEFMRFKVHMEG
SVNGHEFEIEGEGEGRPYEAFQTAKLKVTKGGPLPFTWDILSPQFMYGSKVYIKHPADIPDY
FKLSFPEGFRWERVMIFEDGGIIHVNQDSSLQDGVFIYKVKLRGTNFPSDGPVMQKKTMGL
EACEERMYPEDGALKSEYKEWLKLKDGGHYAAEVKTTYKAKKPVQLPGAYIVDIKLDIVSHN
EDYTIVEQYERAEGRHSTGGMDELYKGGSASQGEELFTGVVPILVELDGDVNGHKFSVRGE
GEGDATIGKLTLKFISTTGKLPVPWPTLVTTLTYGVQCFSRYPDHMKRHDFFKSAMPEGYV
QERTISFKDDGKYKTRAVVKFEGDTLVNRIELKGTDFKEDGNILGHKLEYNFNNPPDSGVRAI
DLPLGENLKARIVQLNEILQVDLEQFQALTDAGVEIGTEVDIINEQGRVVITHNGSSVELIDDLA
HAVRVEKVEG*

>pAZ188_pRSET_10xHis_EK_MDtxRGA_P146 (749 aa)
MRGSHHHHHHHHHHGMASMTGGQQMGRDLYDDDDKDPGRMKDLVDTTEMYLRTIYELE
EEGIVPLRARIAERLEQSGPTVSQTVARMERDGLVHVSPDRSLEMTPEGRSLAIAVMRKHRL
AERLLTDIIGLDIHKVHDEACRWEHVMSDEVERRLVEVLDDVHRSPFGNPIPGLGEIGLDQA
DEPPASHNVYITADKQKNGIKANFTVRHNVEDGSVQLADHYQQNTPIGDGPVLLPDNHYLS
TQTKLSKDPNEKRDHMVLLEFVTAAGITHGMDELYGGTVSKGEENNMAIIKEFMRFKVHME
GSVNGHEFEIEGEGEGRPYEAFQTAKLKVTKGGPLPFTWDILSPQFMYGSKVYIKHPADIPD
YFKLSFPEGFRWERVMIFEDGGIIHVNQDSSLQDGVFIYKVKLRGTNFPSDGPVMQKKTMG
LEACEERMYPEDGALKSEYKEWLKLKDGGHYAAEVKTTYKAKKPVQLPGAYIVDIKLDIVSH
NEDYTIVEQYERAEGRHSTGGMDELYKGGSASQGEELFTGVVPILVELDGDVNGHKFSVR
GEGEGDATIGKLTLKFISTTGKLPVPWPTLVTTLTYGVQCFSRYPDHMKRHDFFKSAMPEG
YVQERTISFKDDGKYKTRAVVKFEGDTLVNRIELKGTDFKEDGNILGHKLEYNFNNPDSGVR
AIDLPLGENLKARIVQLNEILQVDLEQFQALTDAGVEIGTEVDIINEQGRVVITHNGSSVELIDD
LAHAVRVEKVEG*

>pAZ189_pRSET_10xHis_EK_MDtxRGA_D147  (749 aa)
MRGSHHHHHHHHHHGMASMTGGQQMGRDLYDDDDKDPGRMKDLVDTTEMYLRTIYELE
EEGIVPLRARIAERLEQSGPTVSQTVARMERDGLVHVSPDRSLEMTPEGRSLAIAVMRKHRL
AERLLTDIIGLDIHKVHDEACRWEHVMSDEVERRLVEVLDDVHRSPFGNPIPGLGEIGLDQA
DEPDPASHNVYITADKQKNGIKANFTVRHNVEDGSVQLADHYQQNTPIGDGPVLLPDNHYL
STQTKLSKDPNEKRDHMVLLEFVTAAGITHGMDELYGGTVSKGEENNMAIIKEFMRFKVHM
EGSVNGHEFEIEGEGEGRPYEAFQTAKLKVTKGGPLPFTWDILSPQFMYGSKVYIKHPADIP
DYFKLSFPEGFRWERVMIFEDGGIIHVNQDSSLQDGVFIYKVKLRGTNFPSDGPVMQKKTM
GLEACEERMYPEDGALKSEYKEWLKLKDGGHYAAEVKTTYKAKKPVQLPGAYIVDIKLDIVS
HNEDYTIVEQYERAEGRHSTGGMDELYKGGSASQGEELFTGVVPILVELDGDVNGHKFSVR
GEGEGDATIGKLTLKFISTTGKLPVPWPTLVTTLTYGVQCFSRYPDHMKRHDFFKSAMPEG
YVQERTISFKDDGKYKTRAVVKFEGDTLVNRIELKGTDFKEDGNILGHKLEYNFNNPSGVRAI
DLPLGENLKARIVQLNEILQVDLEQFQALTDAGVEIGTEVDIINEQGRVVITHNGSSVELIDDLA
HAVRVEKVEG*

>pAZ190_pRSET_10xHis_EK_MDtxRGA_S148  (749 aa)
MRGSHHHHHHHHHHGMASMTGGQQMGRDLYDDDDKDPGRMKDLVDTTEMYLRTIYELE
EEGIVPLRARIAERLEQSGPTVSQTVARMERDGLVHVSPDRSLEMTPEGRSLAIAVMRKHRL
AERLLTDIIGLDIHKVHDEACRWEHVMSDEVERRLVEVLDDVHRSPFGNPIPGLGEIGLDQA
DEPDSPASHNVYITADKQKNGIKANFTVRHNVEDGSVQLADHYQQNTPIGDGPVLLPDNHY
LSTQTKLSKDPNEKRDHMVLLEFVTAAGITHGMDELYGGTVSKGEENNMAIIKEFMRFKVH
MEGSVNGHEFEIEGEGEGRPYEAFQTAKLKVTKGGPLPFTWDILSPQFMYGSKVYIKHPADI
PDYFKLSFPEGFRWERVMIFEDGGIIHVNQDSSLQDGVFIYKVKLRGTNFPSDGPVMQKKT
MGLEACEERMYPEDGALKSEYKEWLKLKDGGHYAAEVKTTYKAKKPVQLPGAYIVDIKLDIV
SHNEDYTIVEQYERAEGRHSTGGMDELYKGGSASQGEELFTGVVPILVELDGDVNGHKFSV
RGEGEGDATIGKLTLKFISTTGKLPVPWPTLVTTLTYGVQCFSRYPDHMKRHDFFKSAMPE
GYVQERTISFKDDGKYKTRAVVKFEGDTLVNRIELKGTDFKEDGNILGHKLEYNFNNPGVRA
IDLPLGENLKARIVQLNEILQVDLEQFQALTDAGVEIGTEVDIINEQGRVVITHNGSSVELIDDL
AHAVRVEKVEG*

>pAZ191_pRSET_10xHis_EK_MDtxRGA_G149  (749 aa) The IronSenseR
MRGSHHHHHHHHHHGMASMTGGQQMGRDLYDDDDKDPGRMKDLVDTTEMYLRTIYELE
EEGIVPLRARIAERLEQSGPTVSQTVARMERDGLVHVSPDRSLEMTPEGRSLAIAVMRKHRL
AERLLTDIIGLDIHKVHDEACRWEHVMSDEVERRLVEVLDDVHRSPFGNPIPGLGEIGLDQA
DEPDSGPASHNVYITADKQKNGIKANFTVRHNVEDGSVQLADHYQQNTPIGDGPVLLPDNH
YLSTQTKLSKDPNEKRDHMVLLEFVTAAGITHGMDELYGGTVSKGEENNMAIIKEFMRFKVH
MEGSVNGHEFEIEGEGEGRPYEAFQTAKLKVTKGGPLPFTWDILSPQFMYGSKVYIKHPADI
PDYFKLSFPEGFRWERVMIFEDGGIIHVNQDSSLQDGVFIYKVKLRGTNFPSDGPVMQKKT
MGLEACEERMYPEDGALKSEYKEWLKLKDGGHYAAEVKTTYKAKKPVQLPGAYIVDIKLDIV
SHNEDYTIVEQYERAEGRHSTGGMDELYKGGSASQGEELFTGVVPILVELDGDVNGHKFSV
RGEGEGDATIGKLTLKFISTTGKLPVPWPTLVTTLTYGVQCFSRYPDHMKRHDFFKSAMPE
GYVQERTISFKDDGKYKTRAVVKFEGDTLVNRIELKGTDFKEDGNILGHKLEYNFNNPVRAID
LPLGENLKARIVQLNEILQVDLEQFQALTDAGVEIGTEVDIINEQGRVVITHNGSSVELIDDLAH
AVRVEKVEG*

>pAZ192_pRSET_10xHis_EK_MDtxRGA_V150  (749 aa)
MRGSHHHHHHHHHHGMASMTGGQQMGRDLYDDDDKDPGRMKDLVDTTEMYLRTIYELE
EEGIVPLRARIAERLEQSGPTVSQTVARMERDGLVHVSPDRSLEMTPEGRSLAIAVMRKHRL

AERLLTDIIGLDIHKVHDEACRWEHVMSDEVERRLVEVLDDVHRSPFGNPIPGLGEIGLDQA
DEPDSGVPASHNVYITADKQKNGIKANFTVRHNVEDGSVQLADHYQQNTPIGDGPVLLPDN
HYLSTQTKLSKDPNEKRDHMVLLEFVTAAGITHGMDELYGGTVSKGEENNMAIIKEFMRFKV
HMEGSVNGHEFEIEGEGEGRPYEAFQTAKLKVTKGGPLPFTWDILSPQFMYGSKVYIKHPA
DIPDYFKLSFPEGFRWERVMIFEDGGIIHVNQDSSLQDGVFIYKVKLRGTNFPSDGPVMQKK
TMGLEACEERMYPEDGALKSEYKEWLKLKDGGHYAAEVKTTYKAKKPVQLPGAYIVDIKLDI
VSHNEDYTIVEQYERAEGRHSTGGMDELYKGGSASQGEELFTGVVPILVELDGDVNGHKFS
VRGEGEGDATIGKLTLKFISTTGKLPVPWPTLVTTLTYGVQCFSRYPDHMKRHDFFKSAMPE
GYVQERTISFKDDGKYKTRAVVKFEGDTLVNRIELKGTDFKEDGNILGHKLEYNFNNPRAIDL
PLGENLKARIVQLNEILQVDLEQFQALTDAGVEIGTEVDIINEQGRVVITHNGSSVELIDDLAH
AVRVEKVEG*

### Supplemental References

1. Belousov, V.V. et al. Genetically encoded fluorescent indicator for intracellular hydrogen peroxide. *Nat. Methods* **3**, 281-286 (2006).
2. Kaczmarski, J.A., Mitchell, J.A., Spence, M.A., Vongsouthi, V. & Jackson, C.J. Structural and evolutionary approaches to the design and optimization of fluorescence-based small molecule biosensors. *Curr Opin Struct Biol* **57**, 31-38 (2019).
3. Ejike, J.O. et al. A Monochromatically Excitable Green-Red Dual-Fluorophore Fusion Incorporating a New Large Stokes Shift Fluorescent Protein. *Biochemistry* **63**, 171-180 (2024).
4. Pohl, E. et al. Structures of three diphtheria toxin repressor (DtxR) variants with decreased repressor activity. *Acta Crystallogr D Biol Crystallogr* **57**, 619-627 (2001).
5. Ding, X., Zeng, H., Schiering, N., Ringe, D. & Murphy, J.R. Identification of the primary metal ion-activation sites of the diphtheria tox represser by X-ray crystallography and site-directed mutational analysis. *Nature Structural & Molecular Biology* **3**, 382-387 (1996).
6. Tully, M.D. et al. BioSAXS at European Synchrotron Radiation Facility - Extremely Brilliant Source: BM29 with an upgraded source, detector, robot, sample environment, data collection and analysis software. *Journal of synchrotron radiation* **30**, 258-266 (2023).
7. Porod, G. Die Röntgenkleinwinkelstreuung Von Dichtgepackten Kolloiden Systemen - 1 Teil. *Kolloid Z Z Polym* **124**, 83-114 (1951).
8. Fischer, H., Neto, M.D., Napolitano, H.B., Polikarpov, I. & Craievich, A.F. Determination of the molecular weight of proteins in solution from a single small-angle X-ray scattering measurement on a relative scale. *J Appl Crystallogr* **43**, 101-109 (2010).
9. Rambo, R.P. & Tainer, J.A. Accurate assessment of mass, models and resolution by small-angle scattering. *Nature* **496**, 477-481 (2013).
10. Hajizadeh, N.R., Franke, D., Jeffries, C.M. & Svergun, D.I. Consensus Bayesian assessment of protein molecular mass from solution X-ray scattering data. *Sci Rep* **8**, 7204 (2018).
11. Molodenskiy, D.S., Svergun, D.I. & Kikhney, A.G. Artificial neural networks for solution scattering data analysis. *Structure* **30**, 900-908 e902 (2022).
12. Kikhney, A.G., Borges, C.R., Molodenskiy, D.S., Jeffries, C.M. & Svergun, D.I. SASBDB: Towards an automatically curated and validated repository for biological scattering data. *Protein Sci* **29**, 66-75 (2020).
13. Manalastas-Cantos, K. et al. ATSAS 3.0: expanded functionality and new tools for small-angle scattering data analysis. *J Appl Crystallogr* **54** (2021).
14. Panjkovich, A. & Svergun, D.I. CHROMIXS: automatic and interactive analysis of chromatography-coupled small angle X-ray scattering data. *Bioinformatics* (2017).
15. Konarev, P.V., Volkov, V.V., Sokolova, A.V., Koch, M.H.J. & Svergun, D.I. PRIMUS: a Windows PC-based system for small-angle scattering data analysis. *Journal of Applied Crystallography* **36**, 1277-1282 (2003).
16. Svergun, D.I. Determination of the Regularization Parameter in Indirect-Transform Methods Using Perceptual Criteria. *J Appl Crystallogr* **25**, 495-503 (1992).
17. Svergun, D., Barberato, C. & Koch, M.H.J. CRYSOL - A program to evaluate x-ray solution scattering of biological macromolecules from atomic coordinates. *J Appl Crystallogr* **28**, 768-773 (1995).
18. PyMOL The PyMOL Molecular Graphics System, Version 2.5 Schrödinger, LLC. (2022).

19. Hanahan, D. Studies on transformation of Escherichia coli with plasmids. *J Mol Biol* **166**, 557-580 (1983).
20. Studier, F.W. & Moffatt, B.A. Use of bacteriophage T7 RNA polymerase to direct selective high-level expression of cloned genes. *J Mol Biol* **189**, 113-130 (1986).
21. Boyer, H.W. & Roulland-Dussoix, D. A complementation analysis of the restriction and modification of DNA in Escherichia coli. *J Mol Biol* **41**, 459-472 (1969).
22. Platt, R., Drescher, C., Park, S.K. & Phillips, G.J. Genetic system for reversible integration of DNA constructs and lacZ gene fusions into the Escherichia coli chromosome. *Plasmid* **43**, 12-23 (2000).
23. Nelson, K.E. et al. Complete genome sequence and comparative analysis of the metabolically versatile Pseudomonas putida KT2440. *Environ Microbiol* **4**, 799-808 (2002).
24. Kinoshita, S., Udaka, S. & Shimono, M. Studies on the Amino Acid Fermentation. *The Journal of General and Applied Microbiology* **3**, 193-205 (1957).
25. Wennerhold, J. & Bott, M. The DtxR regulon of Corynebacterium glutamicum. *J Bacteriol* **188**, 2907-2918 (2006).
26. Bakkes, P.J. et al. Improved pEKEx2-derived expression vectors for tightly controlled production of recombinant proteins in Corynebacterium glutamicum. *Plasmid* **112**, 102540 (2020).
27. Figurski, D.H. & Helinski, D.R. Replication of an origin-containing derivative of plasmid RK2 dependent on a plasmid function provided in trans. *Proc Natl Acad Sci U S A* **76**, 1648-1652 (1979).
28. Choi, K.H. & Schweizer, H.P. mini-Tn7 insertion in bacteria with single attTn7 sites: example Pseudomonas aeruginosa. *Nat Protoc* **1**, 153-161 (2006).
29. Martinez-Garcia, E. & de Lorenzo, V. Pseudomonas putida as a synthetic biology chassis and a metabolic engineering platform. *Curr Opin Biotechnol* **85**, 103025 (2024).
30. Volke, D.C., Friis, L., Wirth, N.T., Turlin, J. & Nikel, P.I. Synthetic control of plasmid replication enables target- and self-curing of vectors and expedites genome engineering of Pseudomonas putida. *Metab Eng Commun* **10**, e00126 (2020).
